# Enhancer adoption by an LTR retrotransposon generates viral-like particles causing developmental limb phenotypes

**DOI:** 10.1101/2024.09.13.612906

**Authors:** Juliane Glaser, Giulia Cova, Beatrix Fauler, Cesar A. Prada-Medina, Virginie Stanislas, Mai H.Q. Phan, Robert Schöpflin, Yasmin Aktas, Martin Franke, Guillaume Andrey, Christina Paliou, Verena Laupert, Wing-Lee Chan, Lars Wittler, Thorsten Mielke, Stefan Mundlos

**Affiliations:** RG Development & Disease, Max Planck Institute for Molecular Genetics, Berlin, Germany; Microscopy and Cryo-electron Microscopy Service Group, Max Planck Institute for Molecular Genetics, Berlin, Germany; Department of Computational Molecular Biology, Max Planck Institute for Molecular Genetics, Berlin, Germany; Institute for Medical and Human Genetics, Charité-Universitätsmedizin Berlin, Berlin, Germany; Department of Developmental Genetics, Max Planck Institute for Molecular Genetics, Berlin, Germany; Berlin–Brandenburg Center for Regenerative Therapies, Charité-Universitätsmedizin Berlin, Berlin, Germany; Department of Pathology, New York University School of Medicine, Langone Health Medical Center, New York, NY, 10016, USA; Centro Andaluz de Biología del Desarrollo (CABD), Consejo Superior de Investigaciones Científicas/Universidad Pablo de Olavide, Seville, Spain; Department of Genetic Medicine and Development, Faculty of Medicine, University of Geneva, 1211 Geneva, Switzerland; Institute of Genetics and Genomics in Geneva (iGE3), University of Geneva, 1211 Geneva, Switzerland

## Abstract

Mammalian genomes are scattered with transposable elements (TEs). TEs are epigenetically silenced to prevent harmful effects caused by either global activation leading to genome instability or insertional mutation disturbing gene transcription. However, whether the activation of a single element can contribute to pathological phenotypes without directly affecting gene expression is largely unknown. Here, we show that tissue-specific expression of a TE in the embryo leads to the production of viral-like particles (VLPs) which can affect organ formation. Failure to silence an LTR retrotransposon inserted upstream of the *Fgf8* gene results in its co-expression with *Fgf8* in the developing embryo. While local gene regulation is unaffected, the LTR retrotransposon participates in chromatin folding at the locus and adopts the expression of the regulatory domain it is located in. This drives the production of VLPs in the *Fgf8*-expressing cells of the developing limb, triggering apoptotic cell death at the time of digit outgrowth and resulting in a limb malformation resembling human ectrodactyly. This phenotype can be rescued by knock-out or knock-in of the retrotransposon causing mutations preventing its full retroviral cycle. Insertion of the same element at other developmental loci faithfully recapitulates expression according to the neighboring regulatory activity. Our findings provide a mechanism by which TE insertion is incorporated into the local genomic regulatory landscape and show how VLP production in post-implantation embryos can interfere with organ formation.

## Introduction

About half of the mammalian genome consists of transposable elements (TEs)^1^. As their name indicates, these genetic elements can mobilize in the genome, causing insertional mutations and genomic instability^2^. To prevent such harmful effects on the host genome, TEs are transcriptionally silenced by DNA methylation and other epigenetic marks in most somatic cells^3^. TE activation often results from the remodeling of the epigenome. This notably occurs in mammals during pre-implantation development at the time of genome-wide epigenetic reprogramming. In this context, several TE families were shown to participate in establishing pluripotency and embryonic genome activation required for proper development^4–6^. In pathological conditions, such as cancer, neurological disorders, or aging, remodeling of the epigenome has been associated with aberrant TE activation which may play a role in disease progression^7–10^. Yet, very little is known about how TE activation affects post-implantation development. While this critical period of cellular differentiation and organ formation is not subject to global epigenetic changes, failure to establish epigenetic marks at specific TE sequences could occur.

Precise transcriptional output during organogenesis is largely influenced by promoter-enhancer communication and the organization of the genome in the 3D space^11^. TEs can contain transcription factor binding sites, allowing them to act as alternative promoters or enhancers and thereby influence the expression of nearby genes or modify 3D chromatin architecture^12^. New TE insertion or de-silencing is presumed to be mutagenic by disrupting normal transcription processes (*i*.*e*. splicing, polyadenylation, and disruption of an exon or a regulatory element)^13^. However, how an existing regulatory landscape influences TE expression is unknown. Here, we show that an unmethylated LTR retrotransposon can respond to the regulatory elements surrounding its insertion and this is dictated by the organization of the genome into topologically associating domains (TADs). We demonstrated that the expression of this LTR element in the embryo does not impair transcriptional activation but rather affects the maintenance of a cell population during organ formation.

We focused on an endogenous retrovirus (ERV) element from the MusD (type D-related mouse provirus-like) family, likely originating from an infectious simian type D retrovirus^14^. MusD are evolutionary young elements with about a hundred full-length copies in the mouse genome and are highly polymorphic between mouse strains^15–17^. As for many TEs, their expression is high in the inner cell mass of E3.5 mouse embryos, but MusD elements are also active in post-implantation embryos from E7.5 to E13.5 in specific tissues^18^, making them a good model to study the role of TE during organogenesis. Using the example of the mouse Dactylaplasia limb malformation, we demonstrated that enhancer adoption leads to the expression of a MusD element in an *Fgf8*-like pattern in the apical ectodermal ridge (AER) of the developing limb. This full-length element assembles into viral-like particles (VLPs) during embryogenesis, causing premature cell death of the AER cell population responsible for the strong limb phenotype. Based on these results, we posited that enhancer adoption by LTR elements could be a more common phenomenon responsible for developmental malformation. Consistent with this, we validated this enhancer adoption mechanism at other developmental loci and observed VLP production in the muscle progenitor cells of the developing limb when MusD is located in a different regulatory domain. Our results provide a mechanism by which an LTR retrotransposon can be expressed in a time and tissue-specific manner during embryonic development and how this can have severe consequences on organ formation.

## Results

### Unmethylated MusD insertion at the *Fgf8* locus is required to cause Dactylaplasia phenotype

Dactylaplasia is a genetic limb malformation caused by a spontaneous allele, named *Dac1J*, on mouse chromosome 19^19,20^. The *Dac1J* allele was identified as an LTR retrotransposon from the MusD (type D-related mouse provirus-like) family, inserted in the intergenic region between the *Fbwx4* and *Fgf8* genes^21^ (**Fig. 1a-b**). The 7.4kb long MusD-*Dac1J* sequence contains a retroviral sequence with *gag, pro*, and *pol* genes flanked by identical 5’ and 3’ LTR^21^ but, like other MusD elements, lacks an *env* gene^14^ (**Fig.1a**). Dactylaplasia mutants show a severe ectrodactyly-type limb phenotype characterized by missing central digits in both fore- and hind-paws (**Extended data Fig. 1a**) and the appearance of abnormal nail-like structures (**Extended data Fig. 1a-b**). The phenotype is fully penetrant in homozygotes with all limbs affected (**Extended data Fig. 1c**) while heterozygote mutants show a more variable phenotype with 76.2% (32/42) and 85.7% (36/42) affected forelimbs and hindlimbs, respectively (**Extended data Fig. 1d**). However, Dactylaplasia is subject to polymorphisms between mouse strains (**Fig. 1c)** as the MusD-*Dac1J* 5’LTR promoter exhibits differential DNA methylation in the limb^21^ (**Fig. 1d**). The methylation pattern is constitutive among different germ layers of the embryo and in extra-embryonic tissues (**Extended data Fig. 1e**) which results in Dactylaplasia phenotype in 129sv/s2 but not C57Bl6 mice (**Fig. 1c**). For simplicity, “*Dac1J*” and “*Dac1J*-Bl6” will refer to mutant animals with the *Dac1J* insertion in the 129sv/s2 (exhibiting a phenotype) and the C57Bl6 (no phenotype) background, respectively.

**Fig. 1.**
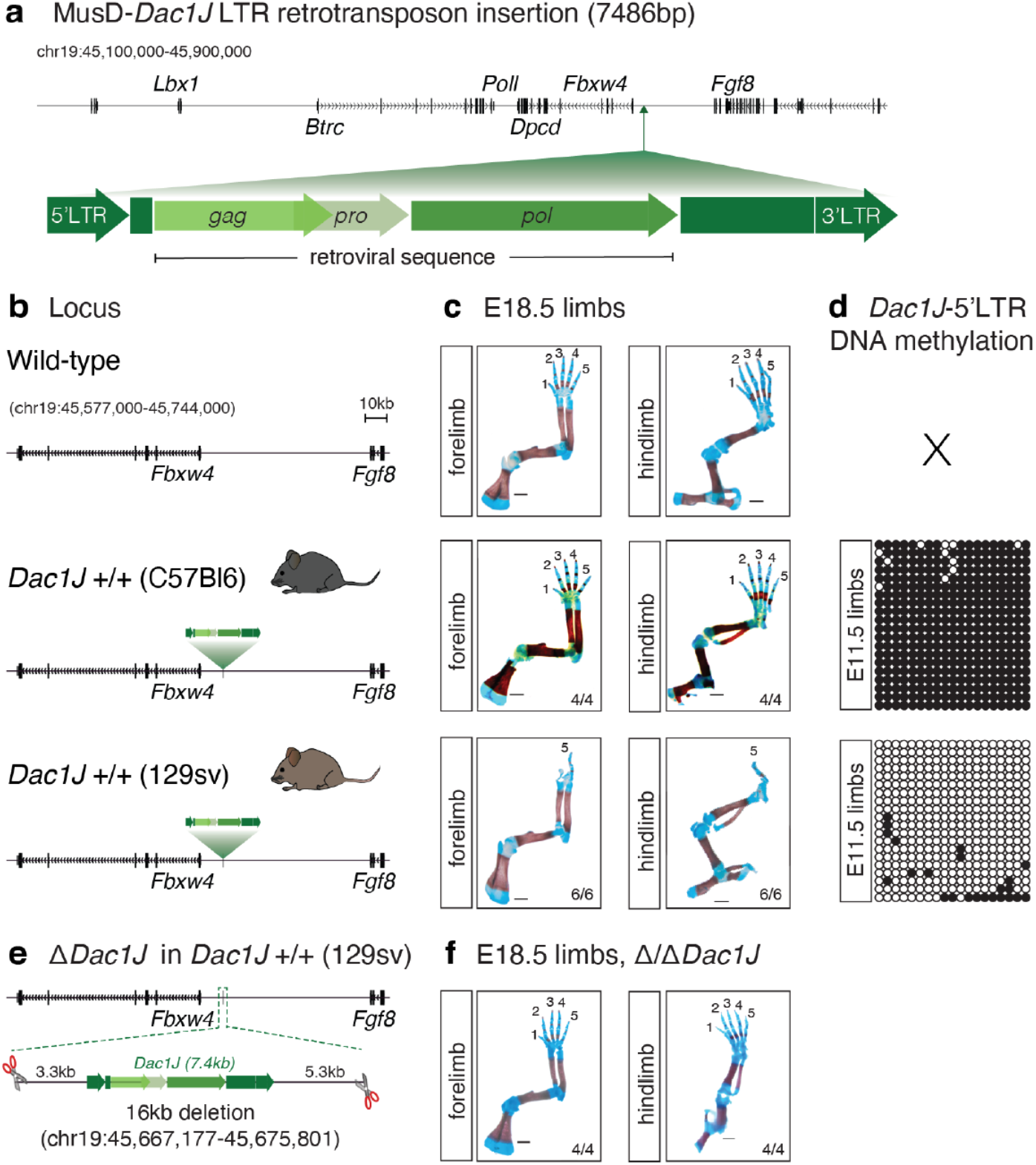
Dactylaplasia limb malformation is caused by unmethylated MusD insertion at the *Fgf8* locus. **a**, Scheme of the 7486 bp MusD-*Dac1J* inserted at the *Fgf8* locus (*mm10, chr19: 45,100,000-45,900,000*). The full-length element contains a retroviral sequence (*gag, pro, pol*) flanked by 5’LTR and 3’LTR. **b**, The mouse *Fbxw4-Fgf8* locus is shown in wild-type (top) or with the intergenic MusD-*Dac1J* insertion in *Dac1J* +/+ C57Bl6 (middle) or 129sv (bottom). **c**, Skeletal analysis of E18.5 wild-type (top), *Dac1J* +/+ C57Bl6 (middle) or 129sv (bottom) fore- and hind-limbs stained with alcian blue (cartilage) and alizarin red (bone). Scale bars 1mm, *n*= 4/4 and 6/6 limbs show a similar phenotype. **d**, DNA methylation status of 19 CpG from the 5’LTR (promotor) from the MusD-*Dac1J* insertion at the *Fgf8* locus in C57Bl6 or 129sv mice measured by bisulfite cloning and sequencing from E11.5 limbs. White circles, unmethylated CpGs; black circles, methylated CpGs. **e**, schematic representation of the CRISPR/Cas9 *Dac1J* deletion in the *Dac1J* +/+ (129sv) line. **f**, Skeletal analysis of E18.5 11/11 *Dac1J* (129sv) fore- and hind-limbs showing complete rescue. Scale bars 1mm, *n*= 4/4 limbs show a similar phenotype.

To functionally prove that the Dactylaplasia phenotype was due to the MusD-*Dac1J* insertion at the *Fgf8* locus, we used CRISPR/Cas9 to delete it from the *Dac1J*+/+ line (**Fig.1e**). *Dac1J* E18.5 embryos with a deletion of MusD-*Dac1J* show a complete rescue of the phenotype (**Fig.1f**), confirming the causative nature of the LTR insertion.

### *Fgf8*-expressing cells but not *Fgf8* activation are affected in *Dac1J* +/+

*Fgf8* is expressed in the apical ectodermal ridge (AER) of the developing limb bud starting at E9.5 where it is responsible for growth and patterning^22^. To elucidate whether the MusD insertion affects the activation of *Fgf8* expression during limb development, we performed single-cell RNA-sequencing from wild-type and *Dac1J* +/+ limb buds at E9.5, E10.5, and E11.5 (**Fig.2a**). We annotated 7 major cell types (**Fig.2b**) which were all present at similar proportions between wild-type and *Dac1J* +/+ at E9.5 and E10.5 (**Fig.2c and Extended data Fig. 2a**). However, at E11.5, a depletion of the AER and dorso-ventral ectoderm cells was detected (**Fig.2c and Extended data Fig. 2a**). In total, we identified a loss of 82.6% of ectoderm cells in the *Dac1J* +/+ E11.5 limbs compared to wild-type (3.7% and 0.7% dorso-ventral ectoderm and 0.9% and 0.1% AER cells in wild-type and *Dac1J* +/+, respectively) (**Fig.2c**). We detected *Fgf8* expression specifically in the AER and dorso-ventral ectoderm cells and show it was not affected at E9.5 and E10.5 in *Dac1J* +/+ limbs compared to wild-type (**Fig. 2d and Extended data Fig. 2b**). At E11.5, despite very few AER cells being present in the *Dac1J* +/+ forelimbs, the remaining cells still expressed *Fgf8* (**Extended data Fig. 2b-c**). Whole-mount *in situ* hybridization at E11.5 confirmed that *Fgf8* can still be detected at the periphery of the AER despite a decrease of expression compared to wild-type (**Extended data Fig. 2b**). We examined the expression of other genes transcribed in the AER in *Dac1J* +/+ which showed proper expression until E10.5 then a decrease at E11.5 when the AER cells have been lost (**Extended data Fig. 2d**). Our results indicates that the lack of *Fgf8* at E11.5 (**Extended data Fig. 2b**) is the result of a lack of AER cells rather than a lack of gene expression.

**Fig. 2.**
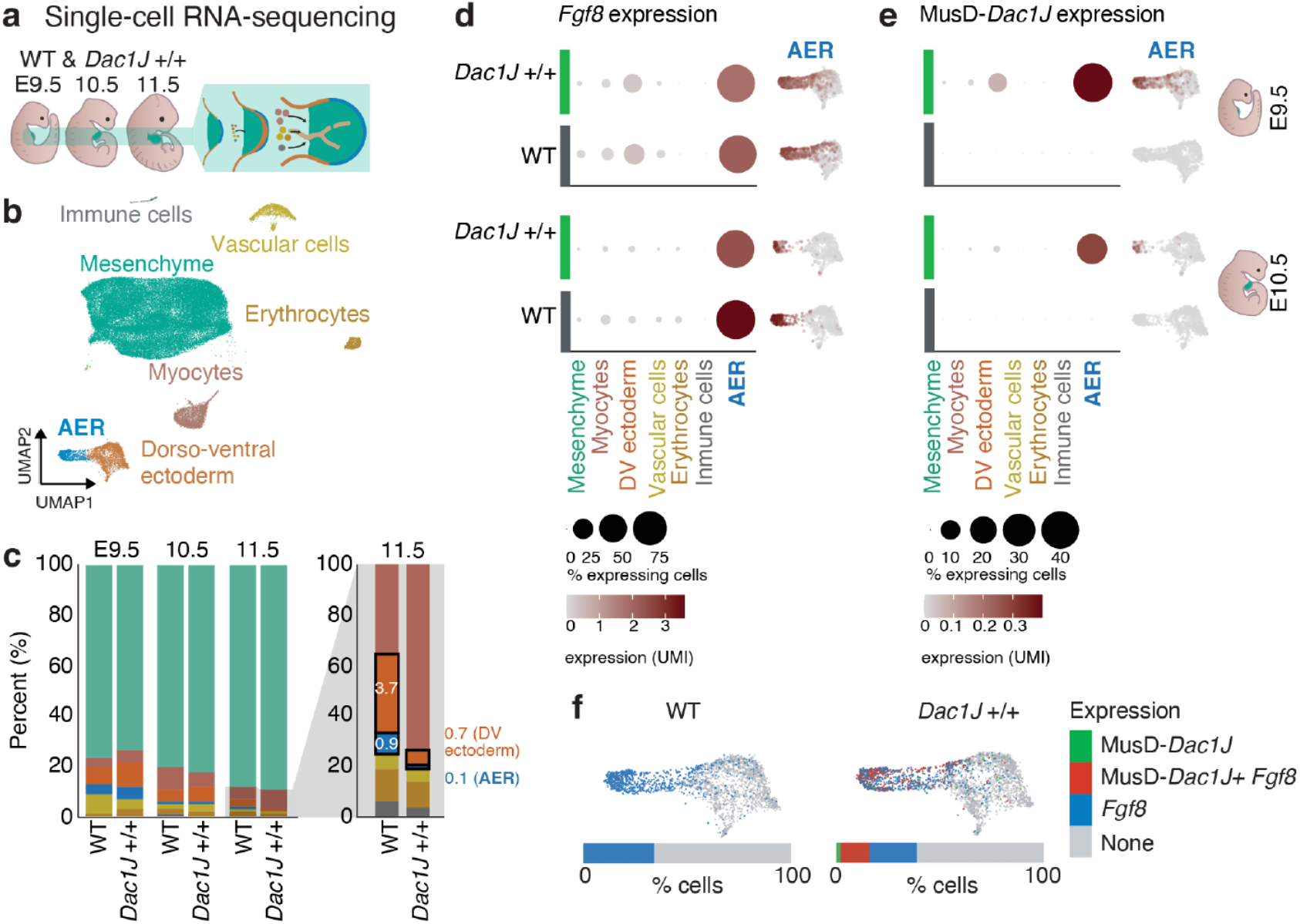
MusD-*Dac1J* transcripts expressed in *Fgf8*-expressing cells before their disappearance. **a**, Schematic representation of the embryos used for limb bud single-cell RNA sequencing. **b**, Uniform manifold approximation and projection (UMAP) showing 7 cell clusters identified via scRNA-seq of E9.5, E10.5, and E11.5 mouse forelimbs from wildtype and *Dac1J* +/+. **c**, Proportion of cell type within the forelimb population in E9.5, E10.5, and E11.5 wild-type and *Dac1J* +/+. Zoom-in view shows a strong decrease of AER and dorso-ventral cells in *Dac1J* +/+ compared to wild-type. **d-e**, dot plots showing expression and percentage of expressing cells for *Fgf8* (**d**) and MusD-*Dac1J* (**e**) in the 7 forelimb cell cluster at E9.5 and E10.5 in wild-type (dark grey) and *Dac1J* +/+ (green). UMI expression in the AER and dorso-ventral ectoderm cell type is also represented next to the respective dot plot for each genotype and stage. **f**, Wild-type (left) and *Dac1J* +/+ (right) AER and dorso-ventral cells colored depending on their status of expressing *Fgf8* and *Dac1J*. Percent of cells expressing either *Dac1J, Dac1J+Fgf8, Fgf8*, or none of them are indicated.

### Co-expression of MusD-*Dac1J* and *Fgf8* in the developing limb

To decipher whether other transcriptional changes occurred before the loss of AER cells, we analyzed the expression of other genes at the *Fgf8* locus during limb development. Using bulk RNA-sequencing from E10.25 fore- and hindlimbs, we did not detect any changes between wild-type and *Dac1J* +/+ (**Extended data Fig. 3a**). We have previously shown that ectopic expression of *Lbx1* and *Btrc* is involved in the pathomechanism of the human split-hand-food malformation type 3 (SHFM3)^23^. In *Dac1J* +/+ embryos, these two genes were expressed normally in all cell types from E9.5 to E11.5 (**Extended data Fig. 3b**).

**Fig. 3.**
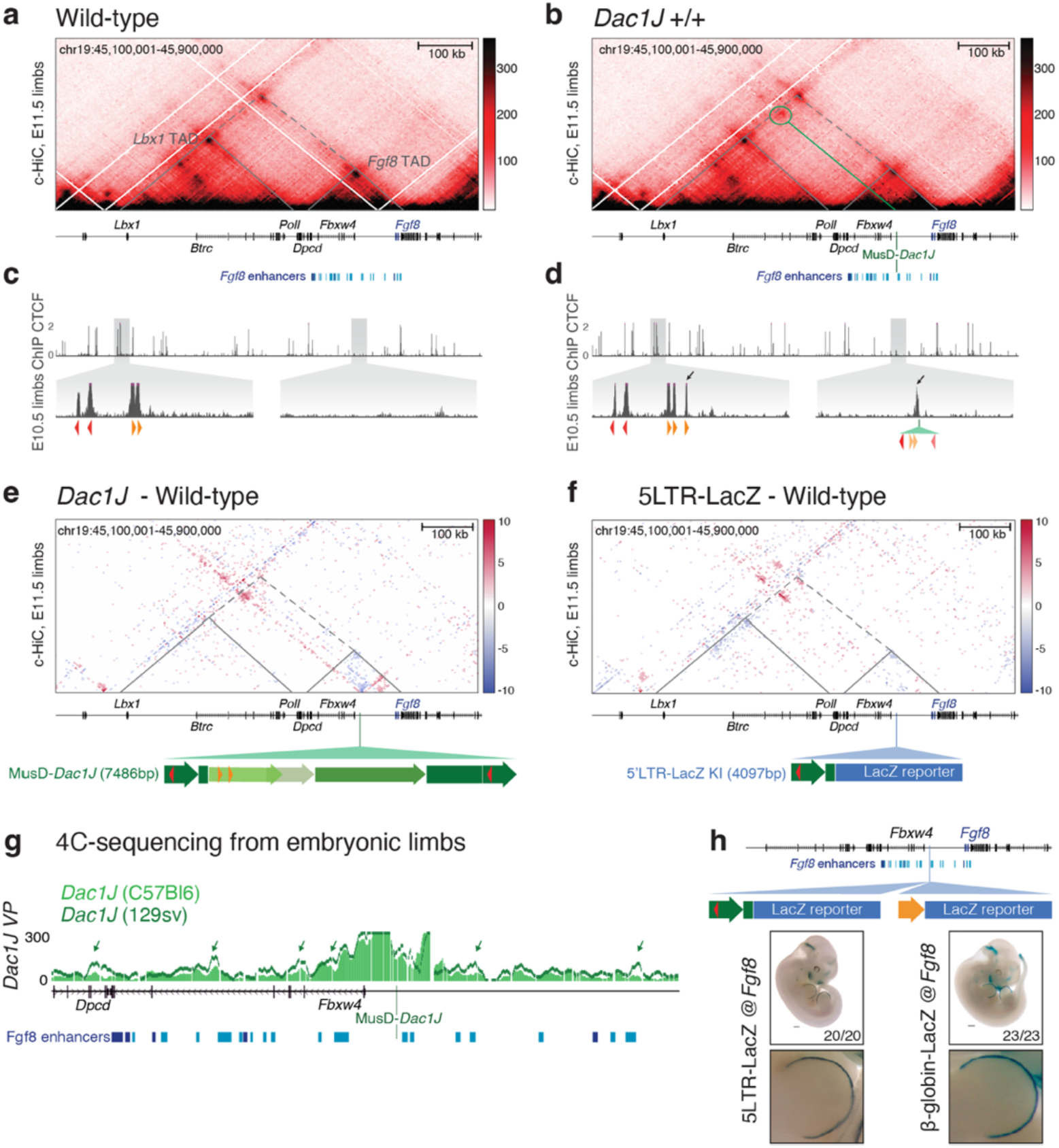
MusD-*Dac1J* insertion leads to ectopic CTCF binding. **a-b**, Capture-Hi-C the *Lbx1/Fgf8* locus (*mm10, chr19: 45,100,000-45,900,000*) from wild-type (**a**) and *Dac1J* +/+ (**b**) E11.5 mouse limb buds. Data are shown as merged signals of *n*=3 biological and 1 technical replicates (**a**) and *n*=2 biological and 2 technical replicates (**b**). The dotted lines indicate the *Lbx1* and *Fgf8* TADs. Published *Fgf8* enhancers are indicated in blue. **c-d**, CTCF ChIP-seq tracks from wild-type (**c**) and *Dac1J* +/+ (**d**) E10.5 mouse limb buds. Zoom-in shows ectopic CTCF peaks in the *Dac1J*+/+ condition (indicated by the black arrow). Orange and red triangles represent CTCF sites in sense and antisense orientation respectively. **e**, Capture-Hi-C subtraction map between wild-type and *Dac1J*+/+. Red and blue colors on the map show gain and loss of interaction due to the MusD-*Dac1J* insertion, respectively. The position of the 7486 bp MusD-*Dac1J* insertion is indicated in green and shows its full-length structure containing the 4 CTCF sites with *p*-value < 10^−4^. **f**, Capture-Hi-C subtraction map between wild-type and *Dac1J* 5LTR-LacZ +/- (*Fgf8* knock-in). The position of the 4097 bp is indicated in blue. **g**, 4C-sequencing with a viewpoint on *Dac1J* showing *Dac1J* +/+ (129sv) and *Dac1J* +/+ (C57Bl6). Green arrows indicate a difference in 4C enrichments. *In vivo*-confirmed *Fgf8* enhancers are indicated in blue, MusD-*Dac1J* insertion point is indicated in green. **h**, Representation of the *Dac1J*-5LTR-LacZ and beta-globin-LacZ knock-ins at the *Fgf8* locus and their beta-galactosidase staining on E11.5 embryos and forelimbs zoom in. Scale bars 500um, *n*= 20/20 and 23/23 embryos show similar staining.

However, when single-cell RNA sequencing reads were mapped to a transcriptome containing the *Dac1J* sequence, we show MusD-*Dac1J* expression in the *Dac1J* +/+ AER cells (**Fig. 2e**). MusD-*Dac1J* transcripts were detected at E9.5 and E10.5 but not at E11.5 when AER cells have vanished (**Fig. 2e and Fig. Extended data Fig. 2c**). Remarkably, nearly all AER cells expressing MusD-*Dac1J* co-expressed *Fgf8* (**Fig. 2f**), showing MusD-*Dac1J* transcription in the exact same pattern as *Fgf8*. Altogether, these results indicate that MusD-*Dac1J* is co-expressed with *Fgf8* in the AER of the developing limb at E9.5 and E10.5 which is followed by a disappearance of most of the AER cells at E11.5.

### MusD-*Dac1J* leads to ectopic CTCF binding within the *Fgf8* TAD

We next asked whether the MusD-*Dac1J* co-expression with *Fgf8* could impact 3D genome architecture. The locus consists of two clearly defined topologically associating domains (TADs)^23^ (**Fig. 3a**). TADs are self-interacting blocks containing genes and their regulatory elements and insulated from each other by the binding of Cohesin and the zinc-finger transcription factor CCCTC-binding factor (CTCF)^24–26^. MusD-*Dac1J* is located in the 175kb size TAD containing *Fbxw4* and *Fgf8* (**Fig. 3b**) as well as previously characterized *Fgf8* enhancers driving its expression in the AER, the hindbrain-midbrain boundary, the branchial arches, and the tailbud (blue bars in **Fig. 3a-b**)^23,27^. Recent data suggest that CTCF binding sites present in retrotransposon sequences can shape the 3D genome organization^28,29^. We thus subjected the MusD-*Dac1J* sequence to CTCF motif scanning and identified several binding sites (red and orange triangles in **Extended data Fig. 4a**). To investigate whether these CTCF sites affect the 3D conformation at the locus, we performed capture-HiC (cHi-C) and CTCF ChIP-sequencing from *Dac1J, Dac1J*-Bl6, and wild-type embryonic limbs. In the *Dac1J* mutants, the two TADs were maintained at the locus but an ectopic chromatin loop was formed by the binding of CTCF to the MusD-*Dac1J* sequence (**Fig. 3a-d and Extended data Fig. 4e**). We also observed increased insulation within the *Fgf8* TAD (**Fig. 3e and Extended data Fig. 4b**). The *Dac1J*-Bl6 embryos do not exhibit any phenotype since the MusD-*Dac1J* promoter is hypermethylated (**Fig. 1c-d**). In these embryos, we did not observe any change in the chromatin conformation or CTCF binding (**Extended data Fig. 4c-e**), showing that the binding of CTCF was sensitive to DNA methylation as shown in other contexts^30^.

**Fig. 4.**
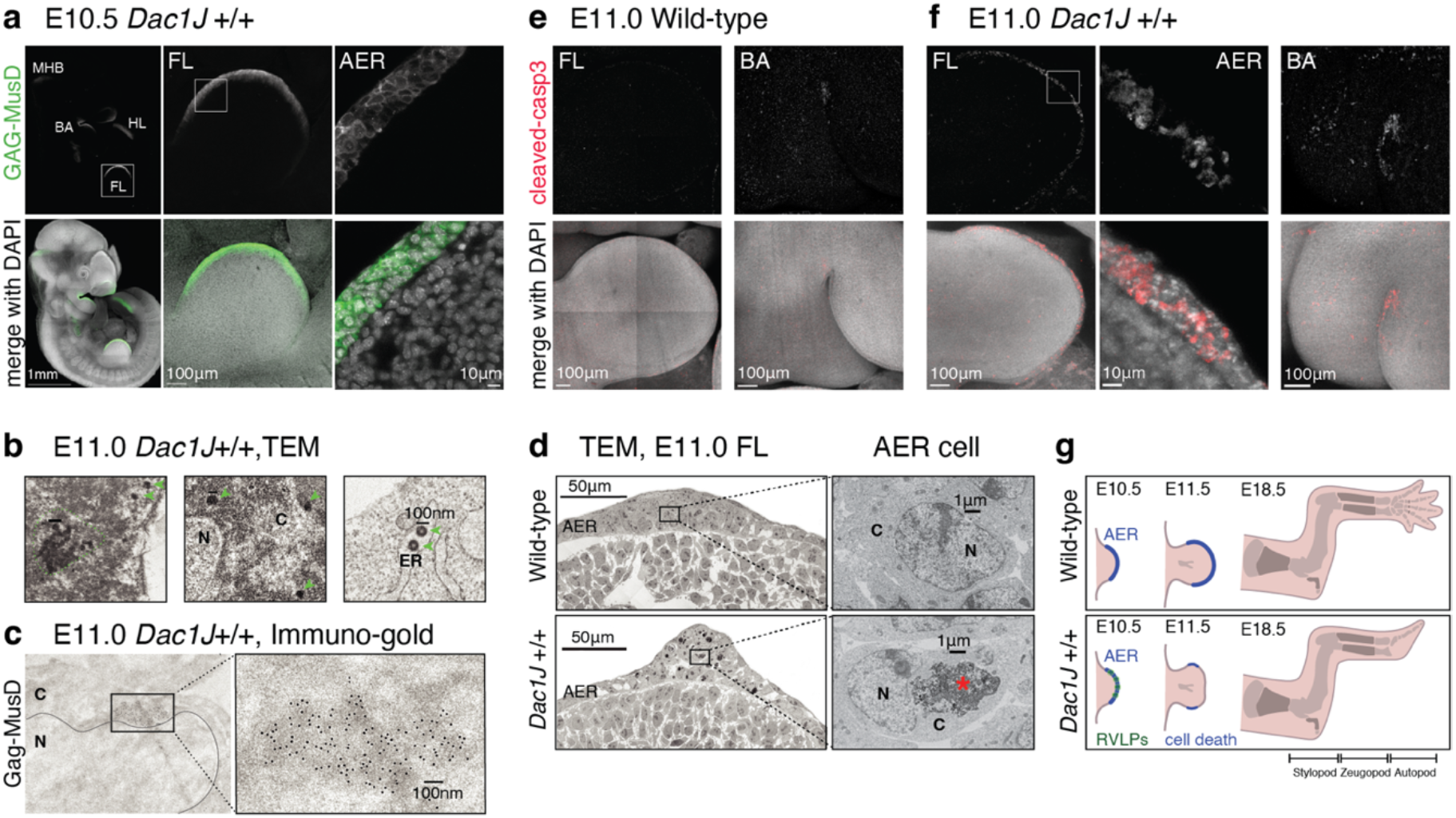
Presence of MusD retroviral-like particles is associated with AER cell death. **a**, anti-GAG-MusD whole-mount immuno-fluorescence on E10.5 *Dac1J*+/+ embryos showing whole embryo (left), forelimb (middle), and AER (right). Scale bars 1mm, 100um, and 10um. At least *n*=3 biological replicates were confirmed. FL, forelimb; HL, hindlimb; BA, branchial arches; MHB, midbrain-hindbrain boundary; AER, apical ectodermal ridge. **b**, TEM analysis on E11.0 *Dac1J* +/+ AER cells. 3 different cells are shown. Green arrows and dotted lines indicate single and aggregated VLPs respectively. Scale bar 100nm. *n*=5 biological replicates and *n*=2 technical replicates were used. C, cytoplasm; N, nucleus; ER, endoplasmic reticulum. **c**, TEM analysis after immuno-gold labeling with anti-GAG-MusD antibody on E11.0 *Dac1J*+/+ AER cell shows cytoplasmic aggregates of GAG. Scale bar 100nm. *n*=2 biological replicates and *n*=6 technical replicates were confirmed. **d**, TEM analysis on E11.0 wild-type and *Dac1J* +/+ forelimbs. Zoom-in view (left) shows the typical structure of an AER cell. Red asterisk indicates an apoptotic body. Scale bars 50um, 100um and 1um. **e-f**, anti-Cleaved-Caspase3 whole-mount immuno-fluorescence on E11.0 wild-type (**e**) and *Dac1J* +/+ (**f**) forelimb and branchial arches. Scale bars 1mm, 100um, and 10um. At least *n*=3 biological replicates were confirmed. **g**, Schematic representation of wild-type and *Dac1J* +/+ limbs at E10.5, E11.5, and E18.5. While the AER of the limb bud is not affected in the *Dac1J* +/+ at E10.5, VLPs are expressed in this tissue. By the time of E11.5, most of the AER cells have undergone cell death. Consequently, at E18.5, the autopod of the *Dac1J* +/+ embryos is severely affected by a lack of metacarpal structures but the stylopod and zeugopod have developed similarly to the wild-type.

According to the loop extrusion model, a chromatin loop is formed between two CTCF sites in a convergent orientation^31^. The MusD-*Dac1J* -mediated chromatin loop seems to originate from the binding of CTCF to an endogenous forward site next to *Lbx1* and an ectopic reverse site in the LTR of the MusD-*Dac1J* sequence (**Fig. 3d**). Yet, several other CTCF sites exist in the MusD-*Dac1J* sequence (**Extended data Fig. 4a**). To test whether only one reverse CTCF can recapitulate the ectopic chromatin loop, we used CRISPR/Cas9 to generate a knock-in of the MusD-*Dac1J* LTR at the *Fgf8* locus. We inserted the MusD-*Dac1J* 5’LTR promoter, which contains one reverse CTCF site, and replaced the retroviral sequence with a LacZ reporter gene (5LTR-LacZ-KI) (**Fig. 3f**). When performing c-HiC from this mutant’s embryonic limbs, we observed a similar ectopic loop as in the *Dac1J* embryos (**Fig 3f, and Extended data Fig. 4f-g**) with an unmethylated 5’LTR promoter (**Extended data Fig. 4h**). This shows that the 5’LTR containing CTCF is sufficient to recapitulate the 3D conformation changes mediated by MusD-*Dac1J*. However, this ectopic contact without the full-length MusD-*Dac1J* retroviral sequence did not lead to any phenotype at E18.5 (**Extended data Fig. 4i**).

### MusD-*Dac1J* promoter adopts the regulatory activity of the *Fgf8* TAD

Our data suggests that a MusD-*Dac1J* mediated ectopic chromatin loop at the *Fgf8* domain neither affects gene expression, nor the Dactylaplasia phenotype. We thus wondered whether it could facilitate MusD-*Dac1J* integration into the *Fgf8* TAD and performed 4C-sequencing with the MusD-*Dac1J* sequence as a viewpoint (**Fig. 3g**). We detected more interaction within the region containing *Fgf8* enhancers in the *Dac1J* mutant compared to the *Dac1J*-Bl6 control, suggesting that the MusD-*Dac1J* 5’LTR could be compatible with *Fgf8* enhancer activity (**Fig. 3g**). We took advantage of the 5LTR-LacZ-KI and generated a similar knock-in where that the LacZ reporter gene is driven by a β-globin minimal promoter instead of the 5’LTR (**Fig. 3h**). β-galactosidase staining in E11.5 embryos with either construct inserted in the *Fgf8* TAD both faithfully recapitulated the native *Fgf8* expression (**Fig. 3h**). This indicates that the MusD-*Dac1J* 5’LTR promoter acts as a sensor adopting the expression pattern of the regulatory landscape at the *Fgf8* locus. Altogether, our data demonstrate that MusD-*Dac1J* can 1) create an ectopic loop when embedded within an existing TAD and 2) contact nearby enhancers to adopt the regulatory activity present in the TAD.

## Assembly of viral-like particles in *Dac1J +/+* AER cells

Having shown that MusD-*Dac1J* does not act as a cis-regulatory element influencing gene expression but rather uses the regulatory landscape for its own transcription, we asked whether this is linked to the lack of AER at E11.5. *Dac1J* is a full-length MusD element with intact ORFs for *gag, pro*, and *pol* (**Fig 1a**). At least nine complete MusD copies with full coding potential exist in the mouse genome, including three shown to be autonomous proviruses still functional for retro-transposition^15^. Sequence alignment revealed that the *Dac1J* GAG and POL protein sequences were almost identical to the three autonomous MusD (**Extended data Fig. 5a**), prompting us to examine whether MusD-*Dac1J* produces retroviral proteins in the developing limb. We used a well-characterized Gag-MusD polyclonal antibody^15^ for whole embryo immunofluorescence assay. We detected Gag-MusD in all analyzed *Dac1J* +/+ embryos (E10.5, *n*=5/5 and E11.0, *n*=4/4) but no staining in the *Dac1J*-Bl6 controls (E10.5, *n*=5/5) (**Fig. 4a and Extended data Fig. 5b-d**). Cytoplasmic Gag-MusD was observed in tissues expressing *Fgf8* such as the mid-hindbrain boundary, the branchial arc, and the AER of both fore- and hindlimbs at E10.5 (**Fig. 4a and Extended data Fig. 5c**), but not in the tailbud (**Extended data Fig. 5c**) and persisted in the AER of *Dac1J* E11.0 embryos (**Extended data Fig. 5d**). This prompted us to examine the presence of viral-like particles (VLPs) in the AER cells. Remarkably, automated transmission electron microscopy (TEM) revealed the presence of cytoplasmic electron-dense particles of 70-90nm-the reported size for MusD VLPs^32^ (**Fig. 4b**). VLPs were detected both in clusters and as single particles close to the nuclear membrane and in the cisternae of the endoplasmic reticulum (**Fig. 4b**). The presence of MusD-derived particles in the AER of *Dac1J* embryos was supported by immuno-gold TEM staining, which detected VLPs at the nuclear membrane labeled by Gag-MusD antibodies in several AER cells (**Fig. 4c and Extended data Fig. 6a-c**). Staining from wild-type control embryos showed no signal (**Extended data Fig. 6d**). This demonstrates the presence of retroviral proteins and particles in the AER cells of the developing *Dac1J* mutant embryos.

**Fig. 5.**
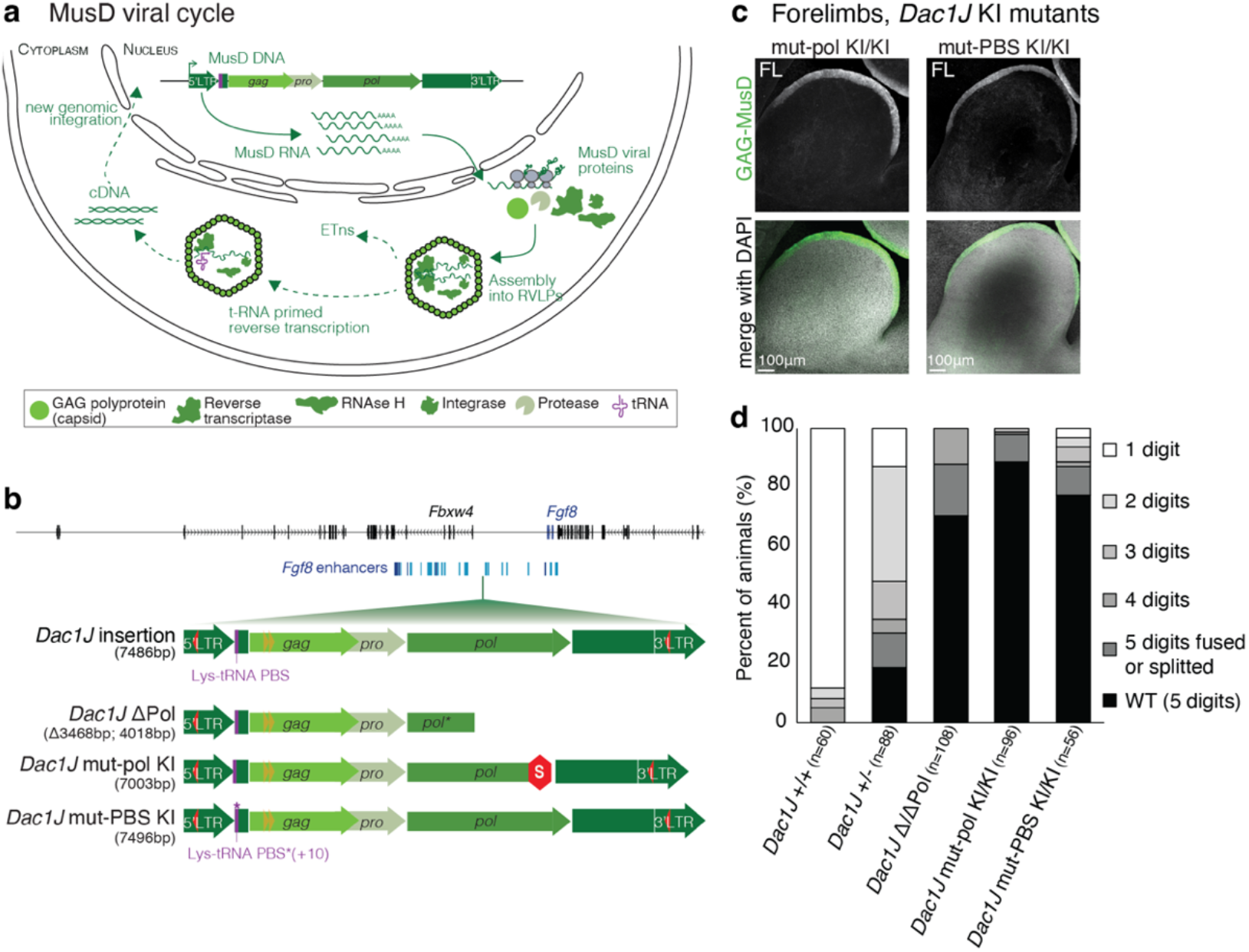
Knock-ins of MusD-*Dac1J* carrying mutation partially rescue Dactylaplasia. **a**, Schematic representation of the MusD viral cycle in a cell. MusD DNA insertion is transcribed into mRNA in the nucleus. MusD RNA is then translated into the Gag, Pro, and Pol retroviral-like proteins in the cytoplasm and assembled into viral-like particles (VLPs). The assembled VLPs contain reverse transcriptase, integrase, and RNAse H and are formed by the Gag capsid which was cleaved by protease. The VLP has then the ability to replicate the non-autonomous ETn retrotransposons but also to undergo t-RNA primed reverse transcription to reintegrate in the genome via a cDNA intermediate (as indicated with the dotted arrows). **b**, Representation of the *Dac1J* insertion along with the 3 mutated versions. **c**, anti-GAG-MusD whole-mount immuno-fluorescence on E10.5 Δ/ΔPol-*Dac1J* (left), mut-pol KI/KI (middle) and mut-PBS KI/KI (right) forelimb. Scale bars 100um. At least *n*=3 biological replicates were confirmed. FL, forelimb. **d**, Histogram of the percentage of animals showing the 6 possible digit phenotype situation in all the different mutants. *n* represents the number of limbs (fore- and hindlimbs) analyzed.

**Fig. 6.**
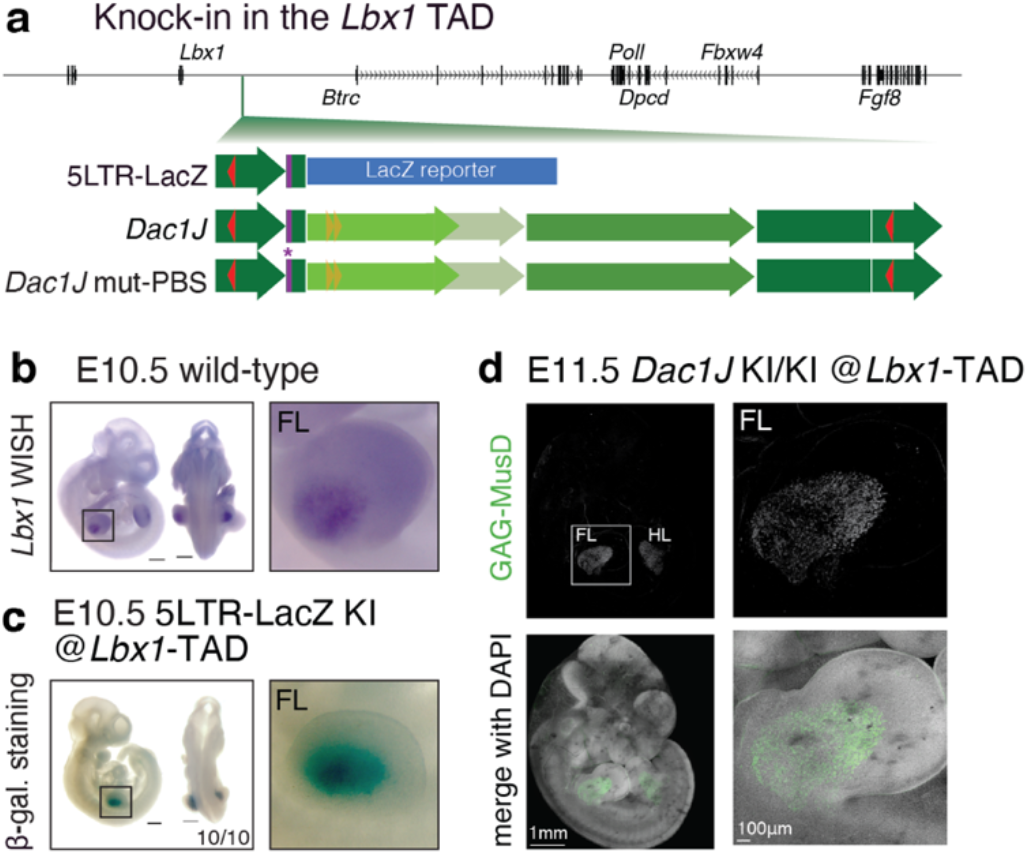
MusD-*Dac1J* adopts the muscle progenitor expression when inserted in *Lbx1* TAD. **a**, representation of the three knock-in lines engineered in the *Lbx1* TAD. **b**, In situ hybridization for *Lbx1* in E10.5 wild-type showing whole embryos and forelimb (FL). Scale bars, 500um. At least *n*=3 biological replicates were confirmed. **c**, beta-galactosidase staining on E10.5 *Dac1J*-5LTR-LacZ (*Lbx1* knock-in) showing whole embryos and forelimb (FL). Scale bars 500um, *n*= 10/10 embryos show similar staining. **d**, anti-GAG-MusD whole-mount immuno-fluorescence on E11.5 Dac1J KI/KI (*Lbx1* knock-in) showing whole embryo (left) and forelimb (right).

### MusD viral-like particles in the AER are associated with aberrant cell death

To decipher how the presence of MusD VLPs during limb embryonic development is linked to the disappearance of AER cells and the Dactylaplasia digit phenotype, we examined TEM images of wild-type and *Dac1J* developing limbs at E11.0. Numerous *Dac1J* AER cells contain up to 4 (or occasionally more) apoptotic bodies and phagocytes (**Fig. 4d and Extended data Fig. 6e-f**). Apoptotic cell death was confirmed by the presence of Caspase-3 in its active form using immuno-staining with a cleaved-Caspase-3 antibody which detected signal in almost all AER cells of both fore- and hindlimbs (*n*= 6/6) and lower signal in a few cells of the branchial arches but no staining in wild-type embryos (*n*= 4/4) (**Fig. 4e-f and Extended data Fig. 7a**). We did not detect cleaved-caspase-3 staining at E10.5 (data not shown) when morphology and proportion of AER cells were not affected (**Fig.2c and Extended data Fig. 2b**), suggesting this cell death must be a rapid process occurring strictly between E10.5 and E11.5. Bulk RNA-sequencing from E11.25 forelimbs shows a significant difference (*p-value* < 0.05) for many genes involved in cell death (**Extended data Fig. 7b**) and genes involved in limb patterning (**Extended data Fig. 7c**). Altogether, our results suggest that the presence of VLPs in the AER leads to non-physiological apoptotic cell death during embryonic development (**Fig. 4g**). This reinforces our finding that Dactylaplasia is caused by the loss of the AER cells rather than the misregulation of *Fgf8* expression, explaining the phenotype of normal proximo-distal outgrowth of the stylopod and zeugopod but strong alteration of the autopod growth (**Fig. 4g**).

### Lack of MusD-*Dac1J* reverse transcription partially rescues the Dactylaplasia phenotype

Autonomous endogenous retroviruses such as MusD elements undergo mRNA splicing and translation to produce the group-specific antigen (Gag) that assembles into a capsid polyprotein, the protease (Pro) for maturation of the gene products, and the polymerase (Pol) encoding for reverse transcriptase (RT), RNase H and integrase (IN) for integration into the genome (**Fig. 5a**). The MusD enzymatic machinery is also used by the closely related but non-autonomous ETn elements for their retro-transposition^14^ (**Fig. 5a**). Having shown that MusD-*Dac1J* is transcribed, translated, and produces VLPs in the AER which are associated with cell death, we wondered which step of the MusD viral cycle causes the phenotype. We hypothesized that the production of MusD VLPs could induce cell death of the AER by either a) sensing the capsid-containing Gag polyprotein and retroviral RNA, b) activation and replication of the ETn elements, or c) undergoing its full retroviral life cycle (**Fig. 5a**). To decipher between these three mechanisms, we designed three mutants carrying modification in the MusD-*Dac1J* sequence (**Fig. 5b**). First, we engineered two mutants with similar consequences on the *pol* gene (producing the reverse transcriptase, RNase H, and integrase): 1) “*Dac1J*-ΔPol” was generated from the *Dac1J* +/+ line and contains a deletion of the 3’ part of MusD-*Dac1J* resulting in a shorter POL protein (**Fig. 5b and Extended data Fig. 8a**) and 2)“*Dac1J-*mut-pol” is a knock-in of the MusD-*Dac1J* with an out-of-frame 493bp deletion in the *pol* gene introducing a 2bp frameshift resulting in a premature STOP codon as previously generated by *Ribet et al*.^15^ (**Fig. 5b**). We verified that the GAG protein was correctly translated in this *Dac1J-*mut-pol by staining with an anti-Gag-MusD antibody, showing that this knock-in recapitulates the same transcription and translation as for the original *Dac1J* insertion (**Fig. 5c and Extended data Fig. 8b**). The lack of a functional POL protein leads to a major rescue of the Dactylaplasia digit phenotype with a milder, not fully penetrant limb phenotype (**Fig 5d and Extended data Fig. 8c**). 29.6% (32/108) and 11.5% (11/96) of the observed limbs were affected in *Dac1J*-11/11Pol and *Dac1J-*mut-pol KI/KI respectively with none of these animals showing the classic one-digit Dactylaplasia phenotype (**Fig. 5d**). None of the observed *Dac1J-*mut-pol heterozygote animals show a phenotype. We validated that the phenotype of these mutants does not depend on variability at the level of DNA methylation at the 5’LTR promoter (**Extended data Fig. 8d**). These observations suggest that the sole presence of the Gag polyprotein and retroviral RNA are not sufficient to drive a phenotype as severe as in the Dactylaplasia mice.

As LTR elements, MusD and ETn have tRNA primer-binding sites (PBS) to initiate reverse transcription and copy their RNA into DNA to reinsert themselves in the genome^33^ (**Fig. 5a**). The *Dac1J*-ΔPol and *Dac1J-*mut-pol mutants prevented the replication of MusD-*Dac1J* itself but also non-autonomous ETn elements. To decipher whether replication of non-autonomous ETn elements plays a role in the development of the Dactylaplasia phenotype, we engineered a third mutant consisting of a knock-in of the full-length MusD-*Dac1J* with a disrupted PBS sequence (**Fig. 5b and Extended data Fig. 8a**). Coding-competent MusD element with a scrambled PBS was no longer able to prime its own reverse transcription but can still replicate its non-autonomous partner, ETn^34^. As for the other mutants, the *Dac1J-*mut-PBS KI/KI embryos express Gag-MusD protein in the cytoplasm of the AER cells (**Fig. 5c and Extended data Fig. 8b**). Interestingly, they also harbor a similar partial rescue of the Dactylaplasia phenotype with 22.6% (14/62) of observed limbs affected (**Fig. 5d and Extended data Fig. 8c**) with none of the heterozygotes showing a phenotype. Altogether, the partial rescues of MusD-*Dac1J* element with disrupted Pol protein or tRNA binding site suggest that full *Dac1J*-derived MusD VLPs with replication potential are required to cause Dactylaplasia phenotype.

### MusD-*Dac1J* adopts the expression of genes surrounding its insertion

Our results demonstrate the severe consequences of activating an endogenous retrovirus element in a tissue-specific manner during embryonic development. We showed that the adoption of the *Fgf8* regulatory activity surrounding the MusD-*Dac1J* insertion is involved in the underlying mechanism. To decipher if this mechanism was specific to the *Fgf8* regulatory domain or a more general mechanism, we inserted the 5’LTR-LacZ construct into the *Lbx1, Shh*, and *Sox9* regulatory domains (**Fig. 6a and Extended data Fig. 9a-b**). When inserted in the *Lbx1* TAD (**Fig. 6a**), adjacent to but separated from the *Fgf8* domain by a TAD boundary (**Fig. 3a**), LacZ expression recapitulated the *Lbx1* expression specifically in muscle progenitors of the developing limbs^35^ (**Fig. 6b-c**). Similarly, 5’LTR-LacZ insertion in the *Shh* and the *Sox9* TADs sensed the surrounding expression in the developing notochord and the limb chondrocytes, respectively (**Extended data Fig. 9c-f**). Of note, no limb staining was observed in the 5’LTR-LacZ-KI at the *Shh* locus, which we attributed to the position of the knock-in as suggested by data from *Symmons et al*.^36^. Altogether, this reveals new properties of an unmethylated 5’LTR, adopting the regulatory activity present in the TAD where they have been inserted, as novel gene promoters^37^.

To investigate whether the production of Gag-MusD polyprotein is specific to *Fgf8*-expressing cells or could also be produced in a different developmental tissue, we engineered knock-ins of MusD-*Dac1J* in its full-length and scrambled PBS forms in the *Lbx1* TAD (**Fig 6a**). Gag-MusD staining was observed in the *Lbx1*-expressing cells of the developing limb; the muscle progenitor cells (**Fig. 6d and Extended data Fig. 9g**). This suggests that MusD VLPs can assemble in any embryonic tissues when adopting the regulatory information from a developmental gene. While the AER and branchial arches are embryonic epithelium, *Lbx1*-expressing cells are from the muscle lineage. In the *Dac1J* KI/KI in the *Lbx1* TAD, Gag-MusD expression was maintained at E11.5 (**Fig. 6d**) while at this stage most of the AER cells are gone in the *Dac1J* +/+ mutants (**Fig. 2c**). When assessing cell death at E12.5, we did not observe muscle progenitor cells undergoing apoptosis, (**Extended data Fig. 9h**) suggesting that these cells from a muscle lineage tolerated VLP production during development. Overall, our results indicate that the cell death response to MusD VLPs during development can affect embryonic cells differently and highlight that mouse post-implantation development can also proceed with the presence of endogenous retrovirus capsid.

## Discussion

In this study, we uncover that an endogenous retrovirus (ERV) element escaping epigenetic silencing can adopt the regulatory activity of developmental genes, resulting in a time- and tissue-specific expression of retroviral products during organ formation. In the Dactylaplasia mutants, VLP assembly in the AER of the developing limb affects cell survival at a critical time preceding digit formation, leading to a lack of digits in newborn mice. The MusD-*Dac1J* insertion is one of the very few reported mutagenic LTR insertions not located in an intron of a gene^13^. Independently from *Dac1J*, a second MusD insertion (*Dac2J*) was reported in the intron of the *Fbxw4* gene, also causing Dactylaplasia^38^. Since *Dac2J* affects *Fbxw4* transcription and both insertions affect a tissue where *Fgf8* is expressed, Dactylaplasia was believed to be primarily caused by disruption of gene transcription^38–41^.

Here, we found that the *Dac1J*-MusD insertion upstream of the *Fgf8* gene leads to two changes at the chromatin level. It creates an ectopic loop (**Fig. 3b-c**) and it increases insulation within the *Fgf8* TAD (**Extended data Fig. 3b**). However, as shown by bulk and single-cell RNA-sequencing analysis, the changes at the 3D genome organization level do not dramatically influence gene activation at the locus during limb development. The MusD insertion instead adopts local regulatory elements, indicating that the MusD 5’LTR can act as a non-specific promoter without disturbing adjacent gene regulation. This enhancer adoption by the MusD element appears to be restricted by TADs and their boundaries as shown by the insertion in the neighboring *Fgf8* and *Lbx1* TADs which results in an AER or a muscle-specific pattern, respectively. Similar results were obtained at other loci, showing that integration of a TE element into the local genomic regulatory landscape is likely to be a more common mechanism. *Fgf8* conditional inactivation in the limb leads to a milder phenotype than in the Dactylaplasia mutants^42,43^ which resembles more closely the double *Fgf8*; *Fgf4* inactivation^44^. Interestingly, the Dactylaplasia phenotype is more comparable to the outcome of AER removal at stage 25 in chick embryos which corresponds to E11.5 in mice, the time point when AER cells are lost in *Dac1J* +/+ embryos^45^. These observations support our data showing that the morphological changes are due to a loss of a cell during this critical step in development rather than changes in gene expression.

A few ERVs carrying intact ORFs for at least Gag, Pol, and Pro proteins have retained the ability to produce viral-like particles (VLPs). Such particles have been observed in pathological contexts when global epigenetic changes occur, resulting in the derepression of many elements from several families, for example in the developing brain^46^, tumor cells^47^, or age-related senescence^9^. It is unclear whether this cellular effect results from the activation of specific elements or the widespread activation of thousands of them. Here, we report the production of VLPs during organ formation in the mouse embryo as a result of epigenetic desilencing of a single MusD element. MusD retrotransposons are endogenous betaretroviruses that have lost the *env* gene but were shown to form intra-cellularized VLPs *in vitro*^15,32^. We have shown that they can also be produced *in vivo* and that the assembly of MusD VLPs in the AER of the developing limb is associated with apoptotic cell death. The *Dac1J* element shows high sequence similarity with the identified MusD “master” copies that are competent and autonomous for retrotransposition^15^ (**Extended data Fig. 5a**), suggesting that *de novo* integration of the MusD-*Dac1J* element could occur, causing genomic instability. However, detecting those insertions in the developing limb is very challenging as they only occur in the cells expressing MusD-*Dac1J* (i.e. the AER and the dorso-ventral ectoderm) which represent less than 4% of all embryonic limb cells (**Fig. 3c**). These cells also die very rapidly following our observation of the VLPs, making it also challenging to choose a time point where new integrations could be detected. The Dactylaplasia phenotype is variable and not fully penetrant in mice carrying a heterozygous insertion of MusD-*Dac1J* (81.2% limb affected) but all homozygotes display a severe digit phenotype, indicating a dose-dependent effect of the VLPs. In contrast, *Dac1J* mutants carrying mutations in the *pol* gene or the PBS had a very mild, low-penetrant phenotype (less than 30% limb affected in homozygotes). While this suggests that the main trigger of the apoptotic signaling resides in the MusD full viral cycle, it also indicates a more complex pathomechanism. MusD Gag and/or RNA immune sensing might also be involved in the phenotype, hinting at various detrimental roles of MusD activation during development.

Our findings raise the question of whether VLP-mediated cell death could happen in other embryonic tissues than the AER. As a result of MusD*-Dac1J* expression, a few cells in the branchial arches of the *Dac1J* embryos undergo cell death. This does not seem to be sufficient to lead to any apparent morphological phenotype, likely because the few dying cells are rapidly replaced by the highly proliferating surrounding tissue. We also showed that MusD-Gag is produced in the *Lbx1*-expressing muscle progenitor cells of the developing limb when MusD is inserted in the *Lbx1* TAD. Cell death was not detected in those cells which can be explained by cell lineage-specificity or dose-dependent effects. We also cannot exclude that the phenotype only appears a few generations after the first insertion as our other knock-in lines were backcrossed before analysis. MusD-derived VLPs could also be tolerated in the embryo as shown in humans where blastocyst development proceeds with the presence of HERVK particles^48^. There is a growing number of studies showing Gag-derived proteins co-opted from retrotransposons producing VLPs^49–51^. Whether or not VLP production could also be co-opted to benefit mouse post-implantation development is yet to be studied. A recent preprint presents evidence that Gag proteins encoded by ERVs are essential for zebrafish and chicken embryonic development^52^.

Despite applying an entire arsenal of whole genome analysis technologies, identifying the causative variants in patients with congenital malformations remains challenging. This raises the possibility that other mechanisms might be the cause of these conditions. Our data suggest that the lack of epigenetic silencing of a single ERV can result in severe malformations induced by tissue and time-specific expression of the ERV in a pattern of the host gene. Variable DNA methylation states leading to incomplete or absent silencing at a critical locus (metastable epiallele)^53^ might be the cause of such conditions, leaving no trace in the genome. Unlike most human ERVs, HERVK (HML-2) retained copies with intact open reading frames for retroviral proteins and was shown to assemble into VLPs in the human early embryo^48^. Long-interspaced nuclear elements (LINEs) are active in humans and their transcription was shown to be associated with immunogenic effects^54^. This supports the idea that the aberrant activation of HERVK or LINEs in developmental tissues could be involved in congenital disorders. This new mechanism may be the key to solving at least part of the many developmental disorders that so far remain poorly understood.

## Methods

The experiments were not randomized. The investigators were not blinded to allocation during experiments and outcome assessment.

### Mice

All animal procedures were conducted as approved by the local authorities (LAGeSo Berlin) under license numbers G0243/18 and G0176/19. For embryo isolation, mice were sacrificed by cervical dislocation and uteri were dissected in PBS. All animal experiments followed all relevant guidelines and regulations. Mice were housed in a centrally controlled environment with a 12 h light: 12 h dark cycle, temperature of 20–22.2 °C, and humidity of 30– 50%. Routine bedding, food, and water changes were performed. The *Dac1J* (129sv/s2) line was obtained from the Jackson Laboratory and the *Dac1J* (C57Bl6) line was obtained through several backcrosses of the *Dac1J* (129sv/s2) animals to C57Bl6 mice.

### Mouse embryonic stem cell targeting

Culture and genome editing of mouse embryonic stem cells (mESCs) was performed as previously described ^23,55^. G4 (wild-type, 129sv/C57Bl6 hybrid) and *Dac1J*-129sv (derived in-house from mouse blastocysts) cells were used. Briefly, mESCs were seeded on a monolayer of CD1 feeder cells and transfection was performed via FuGENE technology (Promega, Cat. #E5911). 8 μg of each CRISPR construct containing the sgRNA of interest delivered through the pX459 vector and 4 μg of the DNA construct for homology-directed repair (HDR) were used. After 24 h, cells were split onto DR4 Puromycin-resistant feeders, and selection with Puromycin was carried out for two days. Resistant-growing clones were then picked and grown into 96-well plates on CD1 feeders. All clones were genotyped by PCR (**Extended Data Table 1**). Positive clones were expanded, their genotyping further confirmed by Sanger sequencing of PCR products and used for further experiments only if the successful modification could be verified. A list of single guide RNAs (sgRNAs, designed using the Benchling CRISPR guides tool) used for CRISPR/Cas9 genome editing is given in **Extended Data Table 2**. All knock-in DNA vectors used for transfection were cloned using Gibson cloning to assemble the KI DNA and the homology arms targeting the *Fgf8, Lbx1, Shh*, or *Sox9* loci. mm10 coordinates of the homology arms used in the KI constructs can be found in **Extended Data Table 3**. The β-globin-LacZ construct contains the minimal β-globin promoter (57bp) upstream of the LacZ gene (3452bp). The *Dac1J*-mutPBS sequence (7496bp) was synthesized and subsequently cloned into a pUC vector by Genewiz. The 5LTR-LacZ constructs contain the MusD-*Dac1J* 5’LTR (318bp) and 5’UTR (331bp, containing the same insertion in the Lys-tRNA PBS as in *Dac1J*-mutPBS) and the LacZ gene (3452bp).

### Generation of mutant mice

Mice were generated from genome-edited mESCs by diploid or tetraploid aggregation^56^. Confirmed CRISPR/Cas9 mutant cell lines after expansion were seeded on CD1 feeders and grown for 2 days before aggregation. Female mice of the CD1 strain were used as foster mothers. The data collection was performed according to the stage of each sample and investigators were not blind to genotype since mouse breeding and analysis required knowledge about the genotype at hand. For Δ*Dac1J*, β-globin-LacZ KI @*Fgf8*, 5LTR-LacZ KI @*Fgf8*, 5LTR-LacZ KI @*Lbx1*, 5LTR-LacZ KI @*Shh*, 5LTR-LacZ KI @*Sox9*, Δ*Dac1J*-Pol, and *Dac1J*-KI @*Lbx1*, only embryos from tetraploid aggregation, were generated for analysis at different embryonic stages. For *Dac1J*-mutPBS KI @*Fgf8, Dac1J*-mutPBS KI @*Lbx1*, and *Dac1J*-mut-pol KI @*Fgf8*, adult mice were generated by diploid aggregation. Founder animals for each mouse line were backcrossed with 129sv wild-type animals for establishing line stock. The selection of animals for analysis and breeding was random. For all MusD-*Dac1J* knock-in mice, the CpG DNA methylation status of the knocked-in 5LTR was assessed by bisulfite sequencing. Mice were used for the study only if the knocked-in 5LTR was hypomethylated.

### Whole-mount in situ hybridization (WISH)

RNA expression in mouse embryos from wild-type and mutants was assessed by WISH using digoxigenin (DIG)-labeled antisense riboprobes for *Fgf8, Lbx1, Shh, and Sox9* transcribed from linearized gene-specific probes (PCR DIG Probe Synthesis Kit; Roche, Cat. #11636090910). Primers for probe generation are listed in **Extended Data Table 4**. Embryos were collected and fixed overnight in 4% PFA in PBS, then washed twice for 30min in PBS with 0.1% Tween (PBST), dehydrated for 30 min each in 25%, 50%, and 75% methanol in PBST, and stored at −20 °C in 100% methanol. WISH was performed as previously described^23^.

### Beta-galactosidase staining

Embryos were dissected in ice-cold PBS and fixed in 4% PFA in PBS for 20 minutes (E11.5 or younger) or 30 minutes (E12.5) and then washed several times in PBS. Embryos were then washed two times for 15 minutes in wash solution (2 mM MgCl2, 0.02% NP-40, and 0.01% C24H39NaO4 in PBS) and finally incubated at 37 °C in X-gal solution (165 mg/mL K4Fe(CN)63H20,

210 mg/mL K3Fe(CN)6 and 40 mg/ml X-gal diluted 1:50 in wash solution). Staining was assessed regularly and the reaction was stopped when LacZ staining was clearly apparent (between 1h to 12h of staining). Samples were then washed several times with PBS and kept at 4 °C in PBS with a few drops of 4% paraformaldehyde. Embryos were imaged using a Zeiss SteREO Discovery V12 microscope and Leica DFC420 digital camera.

### Skeletal preparation

E18.5 fetuses were sacrificed and stored at -20 °C. On the first day of the staining protocol, E18.5 fetuses were kept in H2O for 1–2 h at room temperature and heat shocked at 65 °C for 1min. The skin, abdominal and thoracic viscera were gently removed using forceps. The fetuses were then fixed in 100% ethanol for at least 24 hours. Afterwards, alcian blue staining solution (150 mg alcian blue 8GX in 80% ethanol and 20% acetic acid) was used to stain the cartilage. After 24 h, fetuses were rinsed and post-fixed in 100% ethanol overnight, followed by 24 hours incubation in 0.2% KOH in H2O for initial clearing. The next day fetuses were incubated in alizarin red (50mg l −1 alizarin red S in 0.2% potassium hydroxide) to stain the bones for 24h. Rinsing and clearing were then carried out for several days using 0.2% KOH. The stained embryos were dissected in 25% glycerol, imaged using a Zeiss SteREO Discovery V12 microscope and Leica DFC420 digital camera, and subsequently stored in 80% glycerol.

### Micro-computed tomography

Autopods of 7-months-old wild type and *Dac1J* +/+ (n = 2 per genotype) were wrapped in plastic film and scanned *ex vivo* using a SkyScan 1172 high-resolution micro-computed tomography system (Brucker microCT) at 10-μm resolution, 80 kV, and 124µA. 3D model reconstruction was done with the SkyScan image analysis software (computed tomography analyser and computed tomography volume) (Brucker microCT).

### DNA methylation analysis

Genomic DNA isolated from embryonic tissue was obtained following overnight lysis at 50 °C (100 mM Tris, pH 8.0, 5 mM EDTA, 200 mM NaCl, 0.2% SDS, and proteinase K). DNA was recovered by standard phenol/chloroform/isoamyl alcohol extraction and resuspended in water. Bisulfite conversion was performed on 500–1,000 ng of DNA using the EpiTect Bisulfite kit (Qiagen). Bisulfite-treated DNA was PCR amplified then either cloned and sequenced, or analyzed by pyrosequencing. For cloning-sequencing, at least 8 clones were Sanger sequenced and analyzed with BiQ Analyzer software^57^. Pyrosequencing was performed on the PyroMark Q24 according to the manufacturer’s instructions, and results were analyzed with the associated software. All bisulfite primers are listed in **Extended Table 2**.

### CTCF binding site motif detection

The FIMO (Find Individual Motif Occurrences), MEME suite 5.5.4 (https://meme-suite.org/meme/tools/fimo) was used to detect CTCF motif in the *Dac1J* sequence (AB305072).

### Capture Hi-C

Capture Hi-C protocol was performed as previously described^23^. Briefly, E11.5 mouse limb buds were prepared in 1x PBS and dissociated with trypsin treatment. 2.5 to 5 × 10^6^ nuclei were used for crosslinking and snap-freeze and stored at -80 °C. Snap-frozen pellets were digested with DpnII, ligated, and de-crosslinked. The final library was checked on agarose gel and subsequently used for capture Hi-C preparation. Libraries were sheared using a Covaris sonicator (duty cycle: 10%; intensity: 5; cycles per burst: 200; time: 6 cycles of 60 s each; set mode: frequency sweeping; temperature: 4–7 °C). Adaptors were added to the sheared DNA and amplified according to Agilent instructions for Illumina sequencing. The library was hybridized to the custom-designed SureSelect beads and indexed for sequencing following Agilent instructions. SureSelect enrichment probes were designed over the genomic interval chr19: 44365510-46325510 (mm10) using the online tool of Agilent: SureDesign (https://earray.chem.agilent.com/suredesign/). Novaseq6000 Illumina technology was used according to the standard protocols and with around 400 million 75 or 100 bp (Novaseq6000) paired-end reads per sample.

### Capture Hi-C data processing

Paired-end fastq reads were first trimmed to 50bp to have the same read length across all sequencing runs before alignment to mm10 as well as the appropriate custom mm10 genomes incorporating either the *Dac1J* or the 5LTR-LacZ insertions. Read mapping and further filtering for each library were carried out separately using the HiCUP pipeline v0.8.3^58^ with Bowtie2 v2.5.0^59^ as the aligner and no size selection or filling. Binned and Knight-Ruiz (KR) normalized contact matrices from merged biological replicates were then generated with Juicer tools v1.22.0^60^ for the region of interest (*chr19: 45,100,000-45,900,000*, MAPQ ≥ 30). Before computing subtraction maps, Hi-C matrices from wild-type samples mapped to the appropriate custom genomes were scaled to half that of corresponding heterozygous mutants (*Dac1J*-Bl6 & 5LTR-LacZ) to account for the wild-type allele. Subtraction between KR-normalized matrices (i.e. mutant - wild-type) was then calculated as previously described^61,62^. Specifically, the coverage of the two maps was equalized before element-wise subtraction and z-scaled within the same subdiagonal. Subtraction and matrices visualization at 5kb resolution were done using a custom Python script via the FANC Python API^63^.

### RNA-sequencing

E10.25 and E11.25 forelimb buds were micro-dissected from wildtype and mutant embryos in cold PBS and immediately snap-frozen for storage at 80°C. Total RNA was extracted using the RNeasy Mini Kit according to the manufacturer’s instructions. Samples were poly-A enriched, prepared into libraries using the Kapa HyperPrep Kit, and sequenced on a Novaseq6000 with 75 bp or 100 bp paired-end reads. RNA-seq experiments were performed in triplicates.

### RNA-sequencing data processing

Reads were mapped to the mouse reference genome (mm10) using the STAR mapper^64^ (splice junctions based on RefSeq; options: – alignIntronMin 20–alignIntronMax 500000– outFilterMultimapNmax 5– outFilterMismatchNmax 10– outFilterMismatchNoverLmax 0.1). Reads were subsequently used for expression analysis via the Cufflinks package^65^ (version 2.2.1; default settings). Transcripts of each sample were assembled using Cufflinks provided with reference gene annotations from Ensembl. The resulting assemblies were then merged via Cuffmerge. Heatmap results were visualized with the R package pheatmap.

### single-cell RNA-sequencing

scRNA-seq experiments were performed in single replicates (except for the E9.5 *Dac1J* +/+ samples for which we had 2 replicates) as previously described^23^. Briefly, E9.5, E10.5, and E11.5 limb buds of wild-type and *Dac1J* +/+ embryos were microdissected in ice-cold PBS. A single-cell suspension was obtained by incubating the tissue for 10 min at 37 °C in 200 μL Gibco trypsin-EDTA 0.05% (Thermo Fisher Scientific, Cat. #25300054) supplemented with 20 μL 5% BSA. Trypsinization was then stopped by adding 400 μL 5% BSA. Cells were then resuspended by pipetting, filtered using a 0.40 μm filter, washed once with 0.04% BSA, centrifuged (5min at 150 x g) and resuspended in 0.04% BSA. The cell count was determined using an automated cell counter (Bio-Rad) and cells were subjected to scRNA-seq (10x Genomics, Chromium Single Cell 3′ v2) aiming for a target cell recovery of up to 10,000 sequenced cells per sequencing library. Single-cell libraries were generated according to the 10x Genomics instructions. Libraries were sequenced with a minimum of 230 million 75 bp paired-end reads according to standard protocols.

### single-cell RNA-sequencing data processing

Computational analysis of the sequenced samples was done with Cell Ranger and the Seurat package v.3 (10x Genomics lnc.). Reads were mapped to the mm10 transcriptome customized with an extra chromosome containing the *Dac1J* sequence. Mapping and preprocessing were done with Cell Ranger default parameters version 3.0.2. Genes expressed in fewer than 10 cells were filtered out. Cells were filtered depending on the level of percentage of UMIs mapping to mitochondrial genes, the number of expressed genes, and library size. Only cells with more than 500 detected genes and less than 15% of mitochondrial UMI counts were considered for downstream analysis. When checking for the presence of confounding factors we identified both cell cycle and sex confounding effects. The presence of a cell-cycle effect was checked by using a principal component analysis (PCA) on a set of G2/M and S phase markers genes. We estimated the phase of each cell by assigning a score based on the cell expression of G2/M and S phase markers using the “CellCycleScoring” Seurat function. We used the difference between G2M and S phase scores as a confounding effect in order to only correct for cell cycle phase among proliferating cells while maintaining the difference between stem and progenitor cells as recommended when studying differentiating processes. UMI counts were normalized using scTransform^66^, regressing out the difference between G2M and S phase score and the percentage of UMIs mapping to *Xist* to correct for the difference of ratio between female and male embryos among experiments. We built a common latent cell representation across samples by integrating the sample-wise top 50 cell principal components based on the tap 1000 highly variable genes using the Seurat CCA method^67^. The top 50 principal components of this joint integrated assay were used for the visualization of the Uniform Manifold Approximation and Projection (UMAP). We clustered cells by first constructing a Shared Nearest Neighbor (SNN) Graph based on the Euclidean distance in the first 20 integrated principal components space using the “FindNeighbors” function with k.param set to 20. Cell clusters were defined using the Louvain algorithm as a modularity optimization technique implemented in the function “FindCluster” with the resolution parameter set to 0.2. Visualization of gene expression was computed after a new scTransform normalization run on the merged raw count assays regressing out for cell cycle and sex effect as previously described. For each cluster, conserved markers between mutant and wild types were identified using the Seurat “FindConservedMarkers” function and were then used for cell-type annotation.

### 4C-seq

Mouse limb micro-dissection, cell-dissociation, cross-linking, and nuclei extraction were performed as described above for capture HiC. 4C library preparation was performed as previously described^68^ with modifications described below. A total of 2 to 5 million cells were used as starting material for all 4C-seq libraries. Experiments for the MusD-*Dac1J* viewpoint in *Dac1J*+/+ (C57Bl6) and *Dac1J*+/+ (129sv) samples (n=3) were performed as singletons. 4C-seq primer sequences are listed in **Extended Table 1**. *DpnII* and *BfaI* were used as primary and secondary restriction enzymes, respectively. Parallel inverse PCR reactions were performed to amplify from a total of 1.6 µg template per 4C library and viewpoint. Final 4C-seq libraries were indexed using TruSeq index primers for Illumina (NEB #E7335S) and NEBNext High-Fidelity 2X PCR Master Mix (NEB). Samples were sequenced with Illumina Hi-Seq technology according to standard protocols.

### 4C-seq data processing

4C-seq processing was performed as previously described^69^. Reads were pre-processed and mapped to the mm10 reference genome using BWA. The viewpoint and adjacent fragments 1.5 kb upstream and downstream were removed and a window of 10 fragments was chosen to normalize the data per million mapped reads (RPM). To visualize the data, we created files in bedGraph track format for the read counts of each fragment or in a specified window of fragments.

### ChIP-sequencing

Embryonic limbs were microdissected in ice-cold 1X PBS and pooled. Limbs were washed once with PBS solution and homogenized in 500 μL collagenase solution (0.1% collagenase type 1a (C9891; Sigma), 0.1% trypsin, 5%FCS or chicken serum in DMEM: Ham’s F-12, 1:1) for ∼15 min in a Thermomixer. Then, samples were transferred in a 50 mL Falcon tube through a 40-μm cell strainer and complemented with 10% FCS/PBS solution. Formaldehyde 37% diluted to a final 1% was used to fix the samples for 10 minutes at RT. 1.425 M glycine was used to quench the fixation. Formaldehyde solution was removed by centrifugation (300 x g, 8 min), and fresh lysis buffer (10 mM Tris, pH 7.5, 10 mM NaCl, 5 mMMgCl2, 0.1 mM EGTA with Protease Inhibitor) was added to isolate the nuclei. The samples were incubated for 10 min on ice, centrifuged for 5 min at 480 x g, washed with PBS solution, and snap-frozen in liquid N2. Chromatin was sonicated in a size range of 200– 500 bp by using the Bioruptor UCD-300 system (Diagenode). CTCF-bound chromatin was immunoprecipitated by using the iDeal Kit for Transcription Factors (C01010055; Diagenode) according to the manufacturer’s instructions. Libraries were prepared by using the Nextera adaptors and were sequenced as single-end 50- or 75-bp reads. CTCF antibody (C15410210; Diagenode).

### ChIP-sequencing data processing

Reads were mapped to the mouse reference genome (mm10) using the STAR mapper ^64^ for mapping to the mm10 genome, SAMtools^70^ for filtering, sorting, and removing duplicates, and deepTools^71^ for generating coverage tracks.

### Whole-mount Immunofluorescence (IF)

Embryos were collected in ice-cold PBS and fixed in 4% PFA for 1 h with rotation at 4°C, washed three times in PBS, and stored at 4 °C in PBS + 0.03% Na Azide until staining was performed. On the first day of the protocol, embryos were washed once with PBS for 5min, permeabilized by three times 20 minutes incubations in 0.5% Triton-X/PBS (PBST), and blocked in 5% horse serum/PBST (blocking solution) overnight at 4°C. Primary antibody incubation was performed in the blocking solution for 72 h at 4°C under gentle rotation. Embryos were then washed three times in blocking solution and three times in PBST and incubated overnight in blocking solution at 4°C with gentle rotation. The next day, the secondary antibody diluted in blocking solution was added and embryos were incubated for 48 h at 4°C with gentle rotation. Afterward, embryos were again washed three times in blocking solution and three times in PBST and incubated overnight in DAPI diluted in PBS (1:5000, Sigma-Aldrich, #D9542) at 4°C with gentle rotation. On the last day, embryos were washed three times for 10 minutes each in PBS and post-fixed in PFA 4% at room temperature for 20 min. Prior to clearing, embryos were washed three times with 0.02 M phosphate buffer (PB, 0.025M NaH_2_PO_4_, and 0.075M Na2HPO4, pH 7.4). The clearing was performed by incubation in RIMS (13% Histodenz (Sigma-Aldrich D2158) in 0.02M PB) at 4 °C for at least one day. Whole-mount embryos were then imaged with a Zeiss LSM880 confocal laser-scanning microscope in fast-with Airyscan mode. The rabbit MusD-Gag antiserum was a gift from Thierry Heidemann’s lab as generated by *Ribet et al. 2007*. α-MusD-Gag (1:1000) was used with goat anti-rabbit Alexa-fluorophore 488 (1:1000, Invitrogen #A11008).α-Cleaved-Caspase 3 (rabbit polyclonal, Cell Signaling Technology Cat#9661, Asp175, 1:400) was used with donkey anti-rabbit Alexa-fluorophore 568 (1:1000, Invitrogen #A110042).

### Lysotracker staining

E12.5 embryos were collected in ice-cold PBS and forelimbs were microdissected. Forelimbs were washed once with PBS and then incubated with a 2μM Lysotracker solution (xx) for 30 minutes at 37°C. After incubation, limbs were washed 3 times with PBS and then fixed overnight in PFA 4%. The limbs were then cleared with RIMS and imaged with a Zeiss LSM880 confocal laser-scanning microscope.

### Transmission electron microscopy (TEM)

E11.0 embryos were collected in ice-cold PBS and fixed in 2% PFA and 2.5% glutaraldehyde in PBS for 8h with rotation at 4°C and then kept in PBS at 4°C overnight. After washing three times in fresh PBS, limbs were dissected and post-fixed in 0.5 % (w/v) osmium tetroxide (Science Services, Munich, Germany) in PBS for 1 h at room temperature and then washed four times in PBS for 20 min. Samples were incubated for 30 min in 100 mM HEPES buffer pH 7.4 containing 0.1 % (w/v) tannic acid (Science Services, Munich, Germany), washed three times for 10 min in distilled water, contrasted in 2 % (w/v) uranyl acetate (Sigma-Aldrich, Merck, Darmstadt, Germany) for 1.5 h at room temperature and washed once in distilled water followed by dehydration through a series of increasing ethanol concentrations (30% for 5 min, 50% for 10 min, 70%, 90% and 96% for 15 min each and finally three times 10 min in 100% ethanol, respectively). Samples were then incubated for 5 min in a 1:1 mixture of propylene oxide (Sigma-Aldrich, Merck, Darmstadt, Germany) and absolute ethanol, followed by 5 min in a 1:1 mixture of propylene oxide (Fluka, Merck, Darmstadt, Germany) and SPURR (Low Viscosity Spurr Kit, Ted Pella, Plano GmbH, Wetzlar, Germany) and finally infiltrated with 100 % SPURR at 4°C overnight. SPURR mixture was renewed twice the following day. For polymerization, limbs were transferred into flat embedding molds, surrounded with fresh resin, and incubated for three days at 60°C. 70 nm sections were cut using a Leica UC7 ultramicrotome equipped with a 3 mm diamond knife (Diatome, Biel, Switzerland) and placed on 3.05 mm Formvar Carbon coated TEM copper slot grids (Plano GmbH, Wetzlar, Germany). Sections were post-contrasted using Uranyless EM Stain and ready-to-use 3% Lead Citrate (both Science Services, Munich, Germany). To visualize retrovirus-like particles in the apical ectodermal ridge, sections were imaged automatically using Leginon^72^ on a Tecnai Spirit transmission electron microscope (FEI) operated at 120 kV, equipped with a 4kx4k F416 CMOS camera (TVIPS). For full limb pictures, acquired images (up to 657 micrographs per section through one limb bud) were then stitched to a single montage using the TrakEM2 plugin implemented in Fiji^73,74^.

### Immuno-TEM

Embryos at E11.0 were collected in ice-cold PBS and fixed in 2% PFA and 0.2% glutaraldehyde in PBS for 1h under rotation at 4°C and washed three times in PBS. Limbs were dissected and stained for 30min in 100 mM HEPES buffer pH 7.4 containing 0.1 % (w/v) tannic acid (Science Services, Munich, Germany), washed three times for 10 min in PBS and dehydrated in a series of increasing ethanol concentrations (30% for 5 min, 50% for 5 min, 70%, 90% and 96% for 10 min each and finally three times 10 min in 100% ethanol, respectively). Samples were then incubated for 30 min in a 1:1 mixture of 100% ethanol and London Resin Gold (Plano GmbH, Wetzlar, Germany) and finally infiltrated with 100 % LR Gold at 4°C overnight. On the following day, a fresh mixture of LR Gold was prepared by adding the accelerator Benzil (Plano GmbH, Wetzlar, Germany) in a concentration of 0,2%. Specimens were incubated for two times 2 hours at room temperature and kept overnight at 4°C remaining in this mixture. Before polymerization, a fresh LR Gold solution containing 0.2% Benzil was renewed for 4 hours on the limbs one day later. Limbs were then transferred into flat embedding molds, surrounded with LR Gold / Benzil mixture, and polymerized under a 100 Watt Black-Ray UV Lamp (Plano GmbH, Wetzlar, Germany) for 2 days at 5°C. 80 nm sections were cut using a Leica UC7 ultramicrotome equipped with a 3mm diamond knife (Diatome, Biel, Switzerland) and placed on 3.05 mm Formvar Carbon coated TEM copper slot grids (Plano GmbH, Wetzlar, Germany). Grids were protected against dehydration and unspecific binding by applying the grid on a drop of Aurion blocking solution for goat gold conjugates (Aurion, Wageningen The Netherlands) for 30 min at room temperature. Slices were then incubated on 10 µL of a 1:500 dilution of the rabbit anti-GagMusD primary antibody in Aurion blocking solution overnight at 4°C in a humid chamber. On the following day, grids were washed four times on buffer containing 20 mM TRIS and 0.9% NaCl. As secondary antibodies, goat anti-rabbit 10nm Gold conjugates (British BioCell, Plano GmbH, Wetzlar, Germany) at of 0.08 µg/mL and incubated for two hours at room temperature. The immune reaction was stopped by four times washing on TRIS / NaCl buffer. After a short dip into double distilled water, grids were dried and kept for subsequent examination on a Tecnai Spirit transmission electron microscope (FEI). To visualize the localization of gold conjugates, limb bud sections were imaged at 6500x nominal magnification applying a defocus of -2 µm with a pixel size of 1,7 nm.

## Data Availability

The data generated in this study can be downloaded in raw and processed forms from the National Center for Biotechnology Information Gene Expression Omnibus (GEO) database and are available under accession code GSE246755.

## Code Availability

All analyses were performed using previously published or developed tools, as indicated in the methods section. No custom code was developed or used.

## Acknowledgment

This work was supported by a grant from the Deutsche Forschungsgemeinschaft (DFG) MU 880/16-1 to S.M. J.G. was supported by the HFSP postdoctoral fellowship (LT000465/2019-L). G.A. is supported by Swiss National Science Foundation grants PP00P3_176802 and PP00P3_210995-6. We thank the lab of Thierry Heidemann for generously giving us the antiserum rabbit MusD-Gag. We thank K. Macura, J. Fielder, C. Franke, N. Michaelis, U. Fisher, A. Stiege for technical support, and T. Aktas for intellectual support. We thank the Schulz lab and I. Dunkel for using their pyrosequencer. We thank T. Aktas and W. Schwarzer for their thesis dissertation work. We would like to thank all the members of the Mundlos laboratory for their continuous support and stimulation. Finally, we thank Daniel Ibrahim, Gabriel Cavalheiro, Natalia Benetti, Konrad Chudzik, Philipp Kurbel, Anna Monaco, and Eric van Leen for their critical reading of the manuscript.

## Authors contribution

J.G. and S.M. conceived the project. J.G. designed the experiments and generated transgenic mouse models with the help of G.C. and Y.A. L.W. performed morula aggregation. J.G. and G.C. performed the WISH experiments and the skeletal preparations. W-L.C. performed the micro-CT. J.G. performed and analyzed the bisulfite-cloning sequencing, pyrosequencing, and bulk RNA sequencing. J.G. and M.F. performed the Capture-HiC and the processing was carried out by M.H.Q.P. and R.S. ChIP-sequencing was performed by C.P and analyzed by J.G. Single-cell RNA-sequencing samples preparation was performed by J.G., G.C., G.A., and M.F. and processed by C.A.P-M and V.S. 4C-sequencing was performed by M.F and analyzed by V.L. J.G. performed the whole mount IF. B.F. and T.M. performed the TEM and Immuno-EM experiments. J.G. and S.M. wrote the manuscript with input from G.A., G.C., and M.F., and the manuscript was approved by all authors.

## Competing interest

The authors declare no competing interests.

**Extended data Fig. 1.**
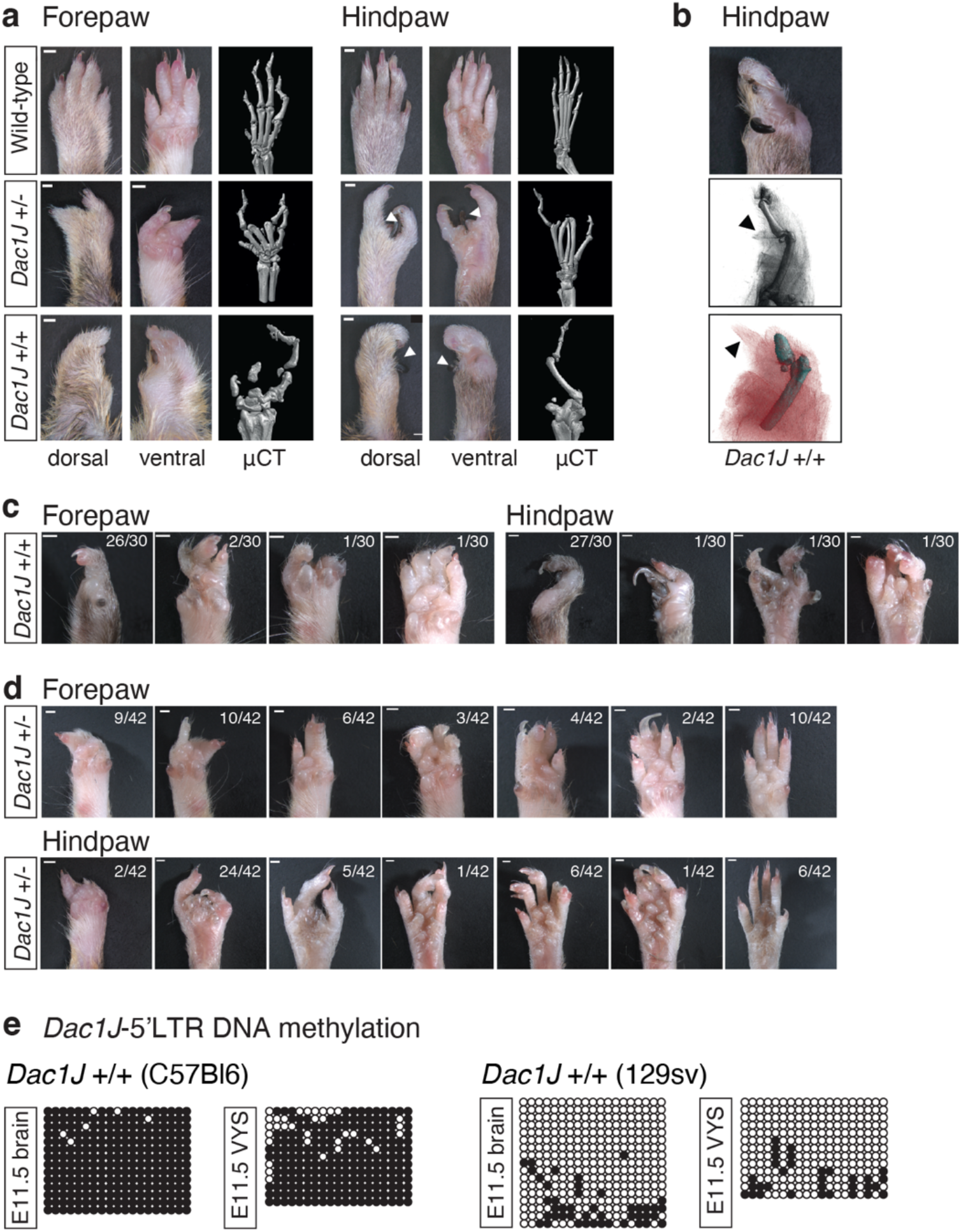
Dactylaplasia phenotype and epigenetic polymorphism. **a**, Dorsal, ventral and micro-computed tomography (µCT) views of wild-type, *Dac1J*+/- (129sv) and *Dac1J*+/+ (129sv) fore- and hind-paws from 7-month-old adults illustrating the phenotype in heterozygotes and homozygotes respectively. Scale bars 1mm, *n*=2 biological replicates were analyzed. White arrows indicate observed nail-like structures. **b**, ventral high-magnification view (top) and micro-computed tomography (µCT) (middle and bottom) of *Dac1J*+/+ (129sv) hind-paw illustrating a typical nail-like-structure (black arrow) which does not contain any bone. **c-d**, Ventral views of the various phenotypes observed in *Dac1J*+/+ (129sv) (**c**) and *Dac1J*+/- (129sv) (**d**) fore- and hindpaws. Scale bars 1mm, n= x/x paws with a similar phenotype. **e**, DNA methylation status of 19 CpG from the 5’LTR (promotor) from the MusD-*Dac1J* insertion at the *Fgf8* locus in a C57Bl6 background (left) or 129sv background (right) measured by bisulfite cloning and sequencing from E11.5 brain, and visceral yolk sac. Constitutive low and high levels of DNA methylation of the MusD-*Dac1J* 5’LTR are observed in the 129sv and a C57Bl6 background respectively. White circles, unmethylated CpGs; black circles, methylated CpGs.

**Extended data Fig. 2.**
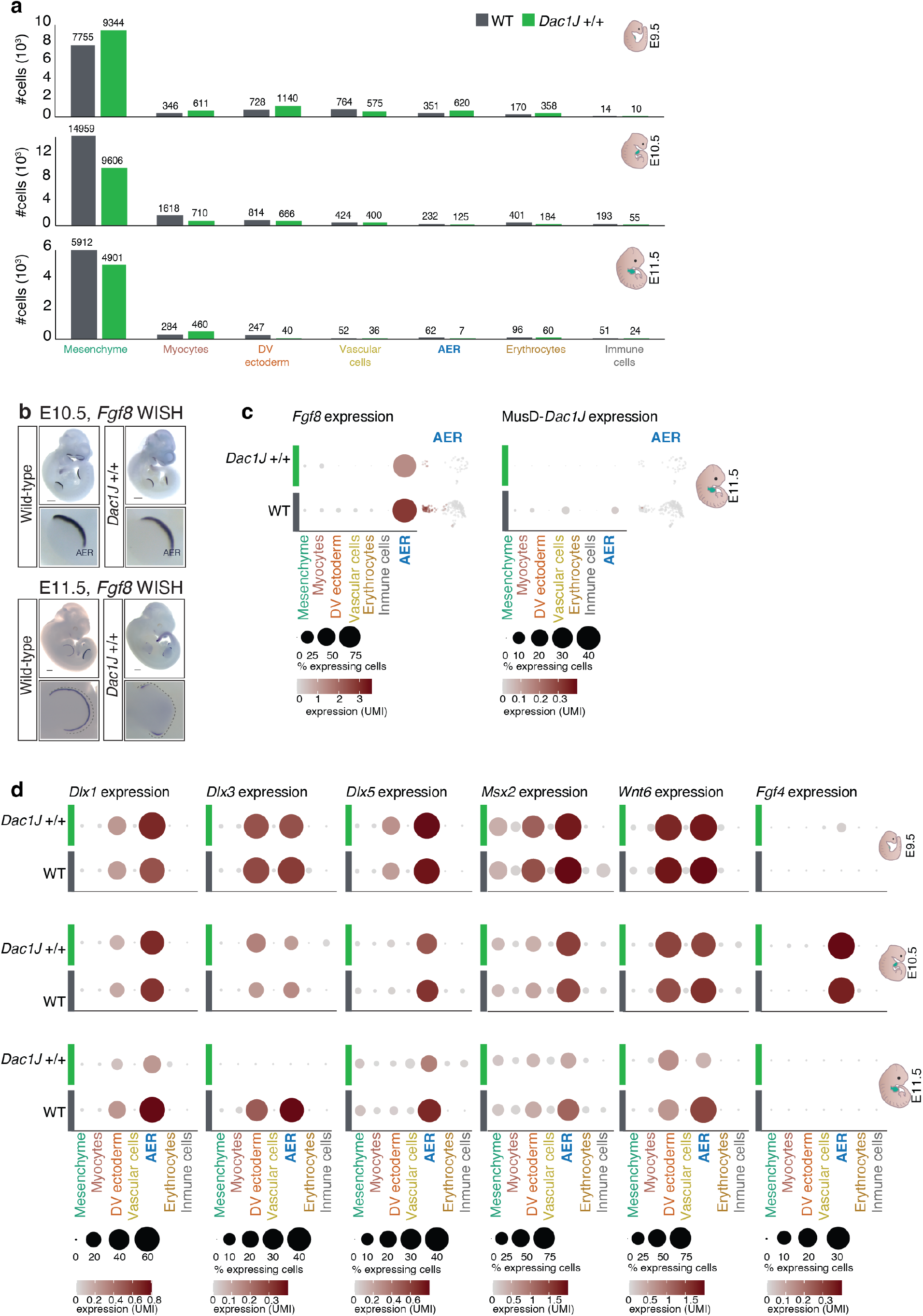
Gene expression changes in the AER of *Dac1J* mutants. **a**, Histogram showing cell number in each cell cluster from wild-type and Dac1J +/+ at E9.5, E10.5, and E11.5. **b**, In situ hybridization for *Fgf8* at E10.5 (top) and in E11.5 (bottom) wild-type and *Dac1J*+/+ showing whole embryos and forelimbs. Dotted lines draw the shape of the forelimb. AER, apical ectoderm ridge. Scale bars 500um, at least *n*= 3 embryos were analyzed per genotype. **c**, Dot plots showing expression and percentage of expressing cells for *Fgf8* and MusD-*Dac1J* in the 7 forelimb cell cluster at E11.5 in wild-type (dark grey) and *Dac1J* +/+ (green) as in **Fig. 2d-e**. UMI expression in the AER and dorso-ventral ectoderm cell type is also represented next to the respective dot plot for each genotype. **d**, dot plots showing expression and percentage of expressing cells for 6 AER genes (*Dlx1, Dlx3, Dlx5, Msx2, Wnt6*, and *Fgf4*) in the 7 forelimb cell cluster at E9.5, E10.5 and E11.5 in wild-type (dark grey) and *Dac1J* +/+ (green).

**Extended data Fig. 3.**
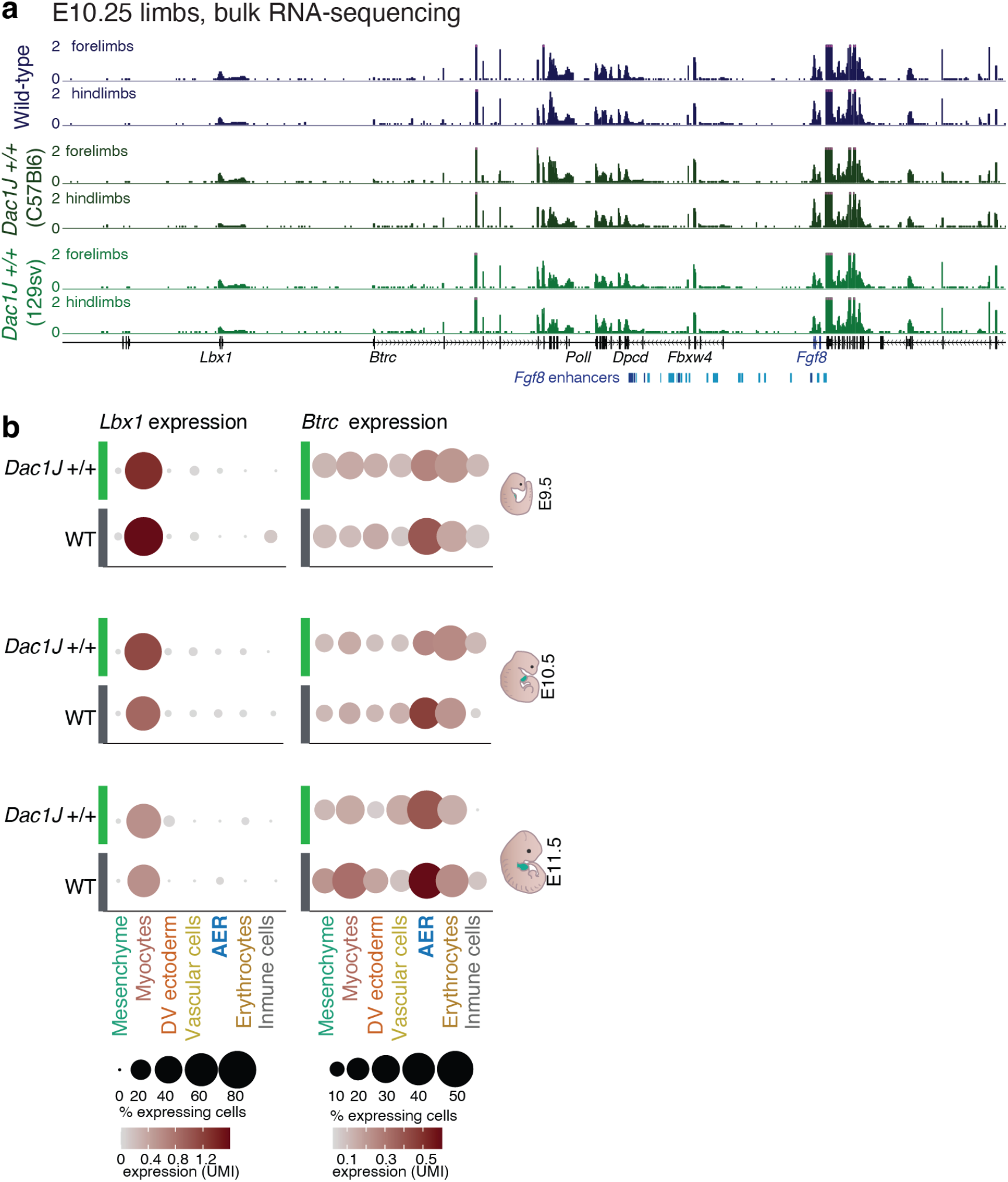
Local gene expression is not impaired in *Dac1J* +/+ embryonic limbs. **a**, Bulk RNA-sequencing tracks from E10.25 fore- and hindlimbs wild type (top), *Dac1J* +/+ (C57Bl6) (middle), and *Dac1J* +/+ (129sv) (bottom). **b**, Dot plots showing expression and percentage of expressing cells for the two genes in the domain adjacent to *Fgf8* (*Lbx1* and *Btrc*) in the 7 forelimb cell cluster at E9.5, E10.5 and E11.5 in wild-type (dark grey) and *Dac1J* +/+ (green).

**Extended data Fig. 4.**
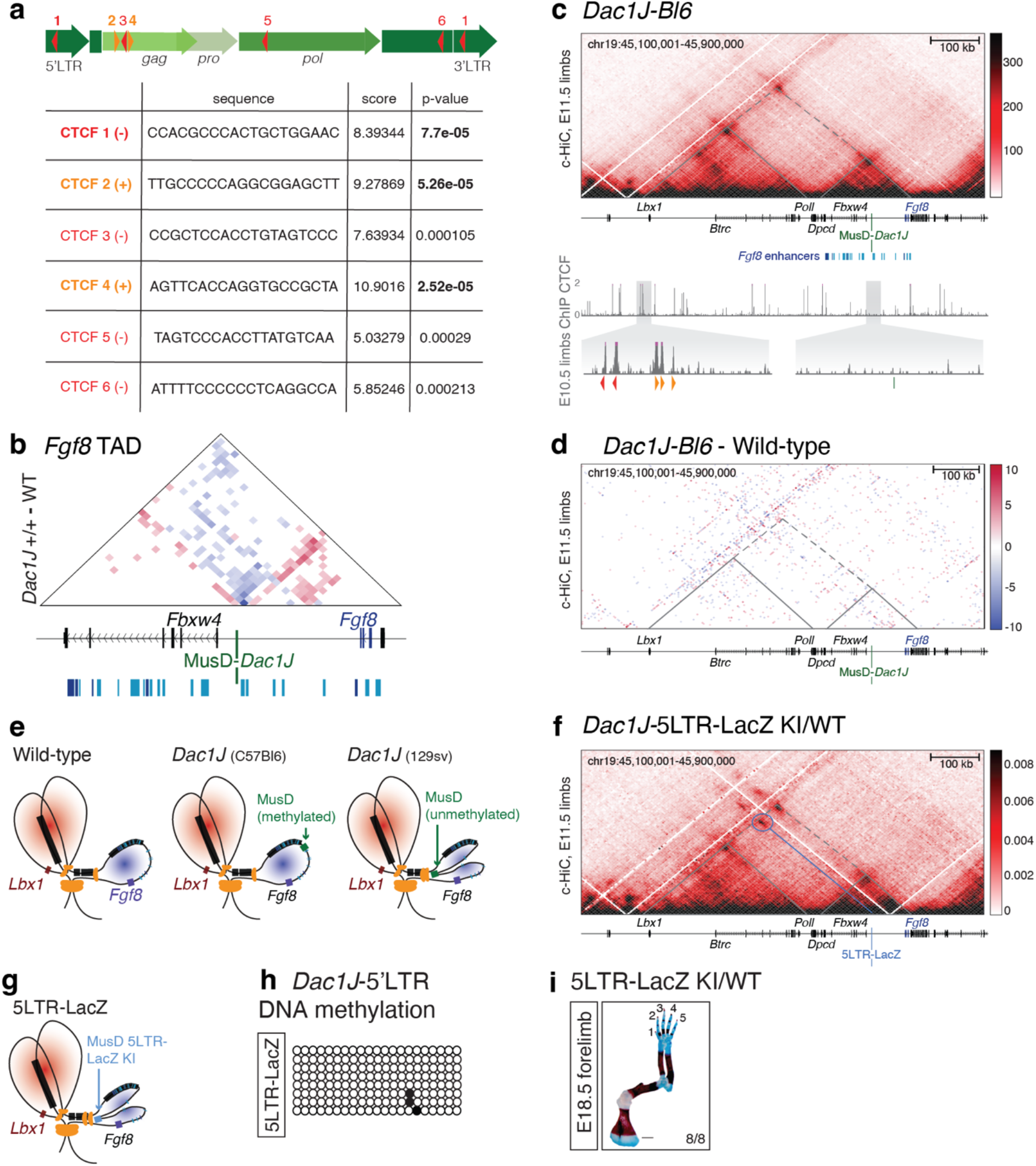
Unmethylated MusD-*Dac1J* insertion induces 3D conformation changes in the *Fgf8* domain. **a**, Sequences of the 6 CTCFs binding sites detected in the MusD-*Dac1J* as identified with the FIMO (Find Individual Motif Occurrences) suite. FIMO score and p-value are indicated for each binding site. Orange and red triangles represent sense and antisense CTCF sites respectively. **b**, Zoom-in from the Capture-Hi-C subtraction map between wild-type and *Dac1J*+/+ (129sv) in **Fig 2e** showing the *Fgf8* TAD. **c**, Capture-Hi-C at the *Lbx1/Fgf8* locus (*mm10, chr19: 45,100,000-45,900,000*) from *Dac1J* +/+ (C57Bl6) from E11.5 mouse limb buds. and ChIP-sequencing tracks from E10.5 mouse limb buds. as in **Fig. 2a-b**. Data are shown as merged signals of *n*=2 biological and 2 technical replicates. **d**, Capture-Hi-C subtraction map between wild-type and *Dac1J*+/+ (C57Bl6) show no differences. **e**, Schematic representation of the 3D conformation at the *Lbx1/Fgf8* locus in wiltype, *Dac1J* +/+ (C57Bl6), and *Dac1J* +/+ (129sv). **f**, Capture-Hi-C at the *Lbx1/Fgf8* locus (*mm10, chr19: 45,100,000-45,900,000*) from *Dac1J* -5LTR-LacZ KI +/-E11.5 mouse limb buds. Data are shown as merged signals of *n* = 3 biological replicates. **g**, Schematic representation of the 3D conformation at the *Lbx1/Fgf8* locus in 5LTR-LacZ KI. **h**, DNA methylation status of 21 CpG from the 5’LTR (promotor) from the MusD-5LTR-LacZ KI at the *Fgf8* locus. **i**, Skeletal analysis of E18.5 5LTR-LacZ KI +/-forelimbs stained with alcian blue (cartilage) and alizarin red (bone). Scale bars 1mm, *n*= 8/8 show a similar phenotype.

**Extended data Fig. 5.**
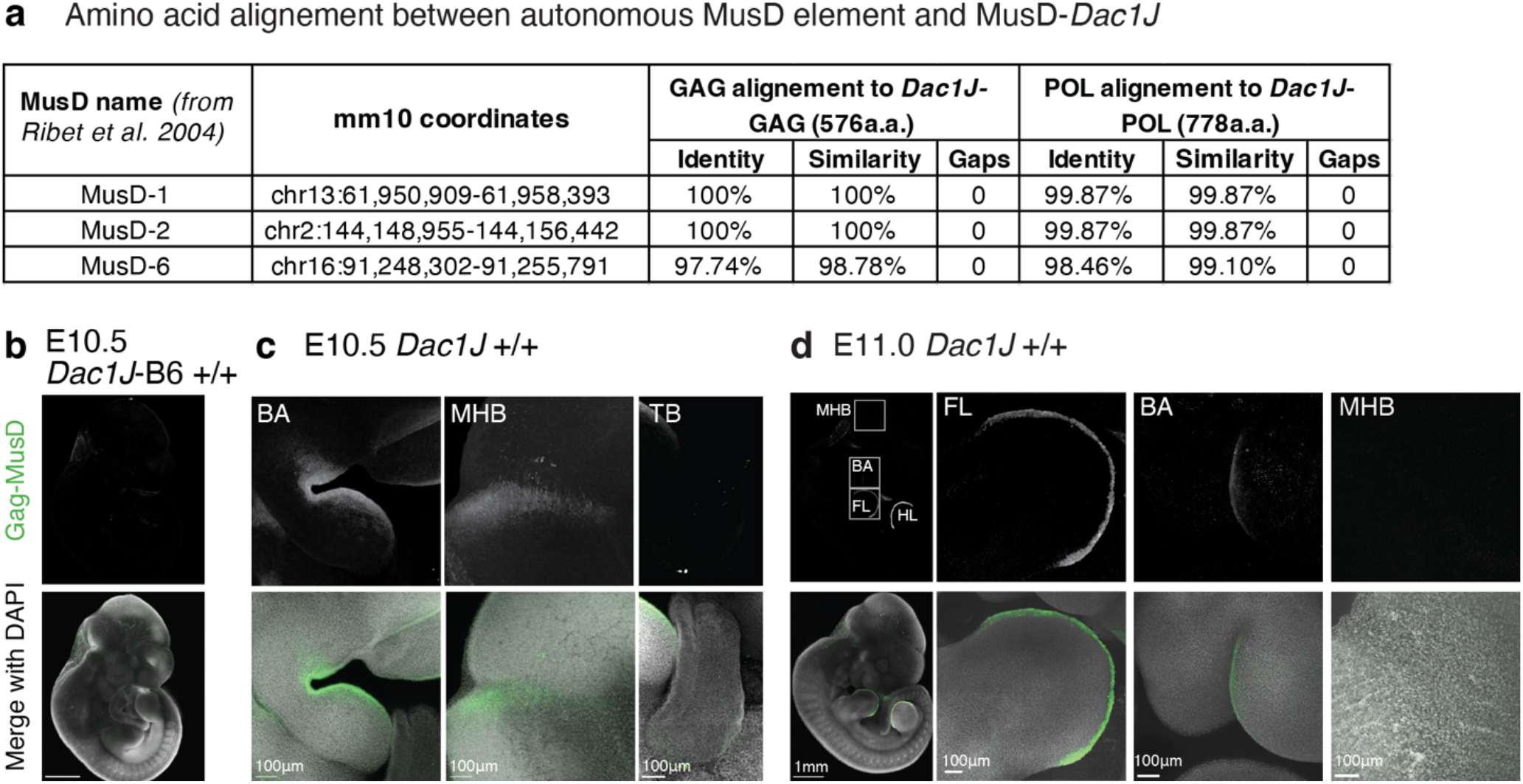
Gag-MusD expression in the *Dac1J* mutant embryos. **a**, Table showing the percentage of amino-acid identity between MusD-Dac1J and the three MusD (MusD-1, -2, and -6) identified as autonomous for retro-transposition in *Ribet et al 2004*. Identity, similarity, and gaps of amino-acids are indicated for both GAG and POL proteins. **b**, anti-GAG-MusD whole-mount immuno-fluorescence on E10.5 Dac1J-Bl6 +/+ embryos showing whole embryo with no staining. Scale bars 1mm. At least n=3 biological replicates were confirmed. **c-d**, anti-GAG-MusD whole-mount immuno-fluorescence on E10.5 (**c**) and E11.0 (**d**) *Dac1J* +/+ embryos showing branchial arches (BA), midbrain-hindbrain boundary (MHB), tailbud (TB), whole embryo and limbs (FL, forelimbs; HL hindlimbs). Scale bar 100nm.

**Extended data Fig. 6.**
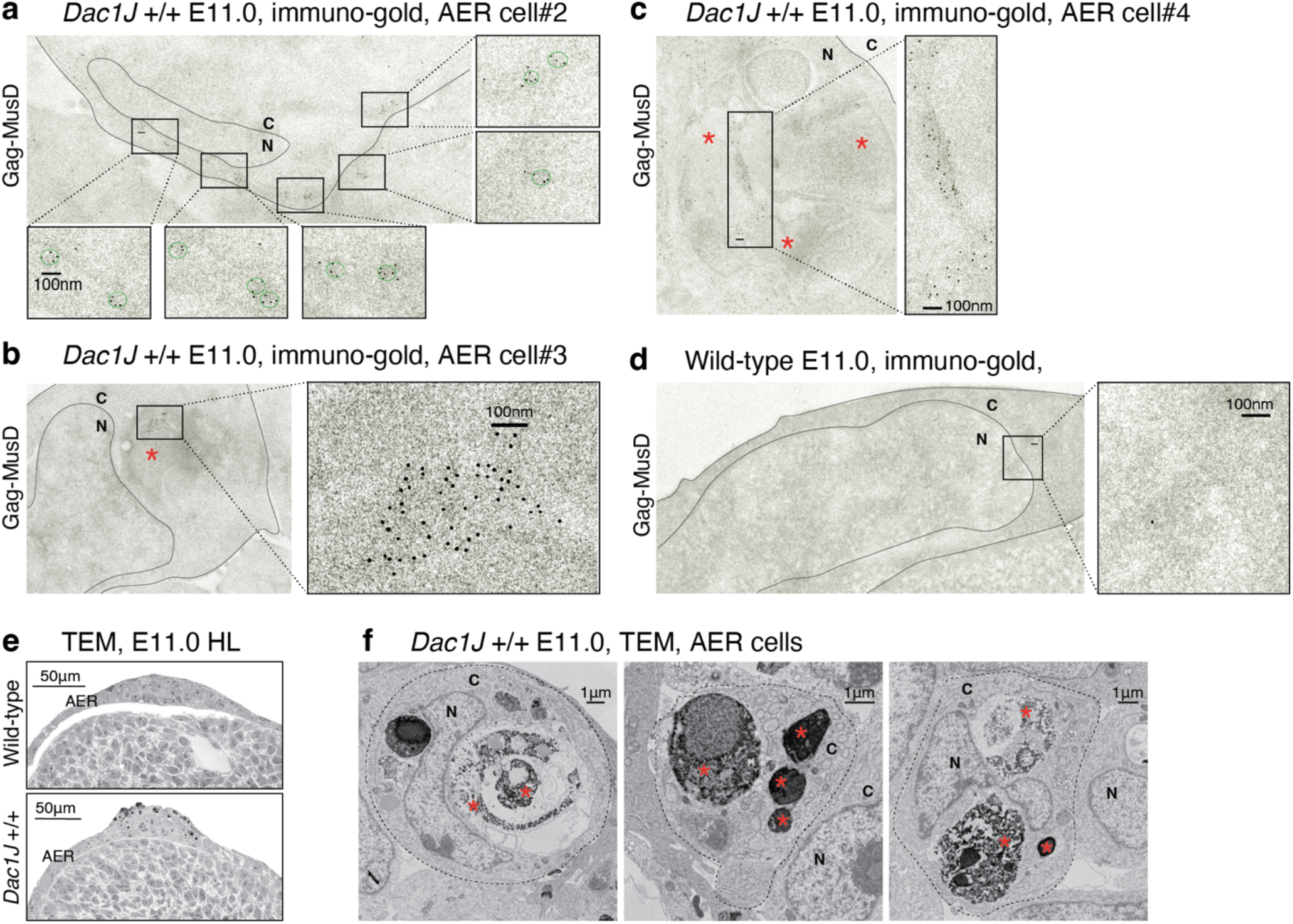
TEM analysis of *Dac1J* embryonic limbs. **a-c**, TEM analysis after immuno-gold labeling with anti-GAG-MusD antibody on E11.0 *Dac1J* +/+ AER cells shows cytoplasmic aggregates of GAG. Scale bar 100nm. *n*=2 biological replicates and *n*=6 technical replicates were confirmed. C, cytoplasm; N, nucleus. Green circle represents one RVLP with Gag capsid. Red Asterix indicates an apoptotic body. **d**, TEM analysis after immuno-gold labeling with anti-GAG-MusD antibody on E11.0 wild-type control AER cell shows no staining. **e-f**, TEM analysis on E11.0 *Dac1J* +/+ hindlimbs (**e**) and zoom-in view on 4 forelimb AER cells showing apoptotic bodies and phagocytes (**f**). Scale bars 50um and 1um.

**Extended data Fig. 7.**
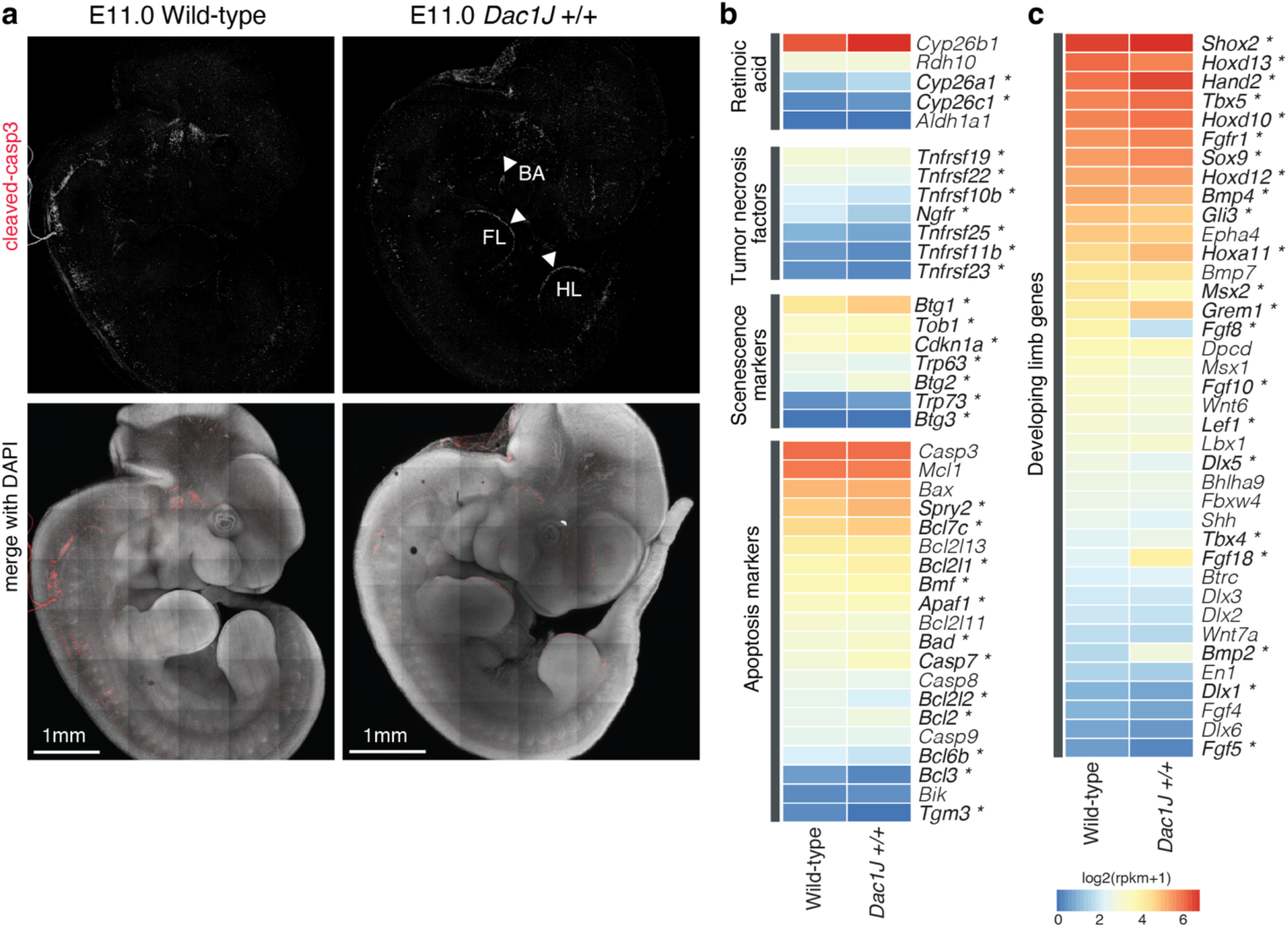
Apoptotic cell death in the *Dac1J* embryos. **a**, anti-Cleaved-Caspase3 whole-mount immuno-fluorescence on E11.25 wild-type and *Dac1J* +/+ showing whole. Scale bars 1mm and 100um. At least *n*=3 biological replicates were confirmed. **b-c**, Heat-map showing log2 (rpkm+1) of selected apoptosis (**b**) and developing limb (**d**) genes. Genes showing significant (p-value < 0.05) expression changes in *Dac1J* +/+ compared to wild-type are indicated with an asterisk.

**Extended data Fig. 8.**
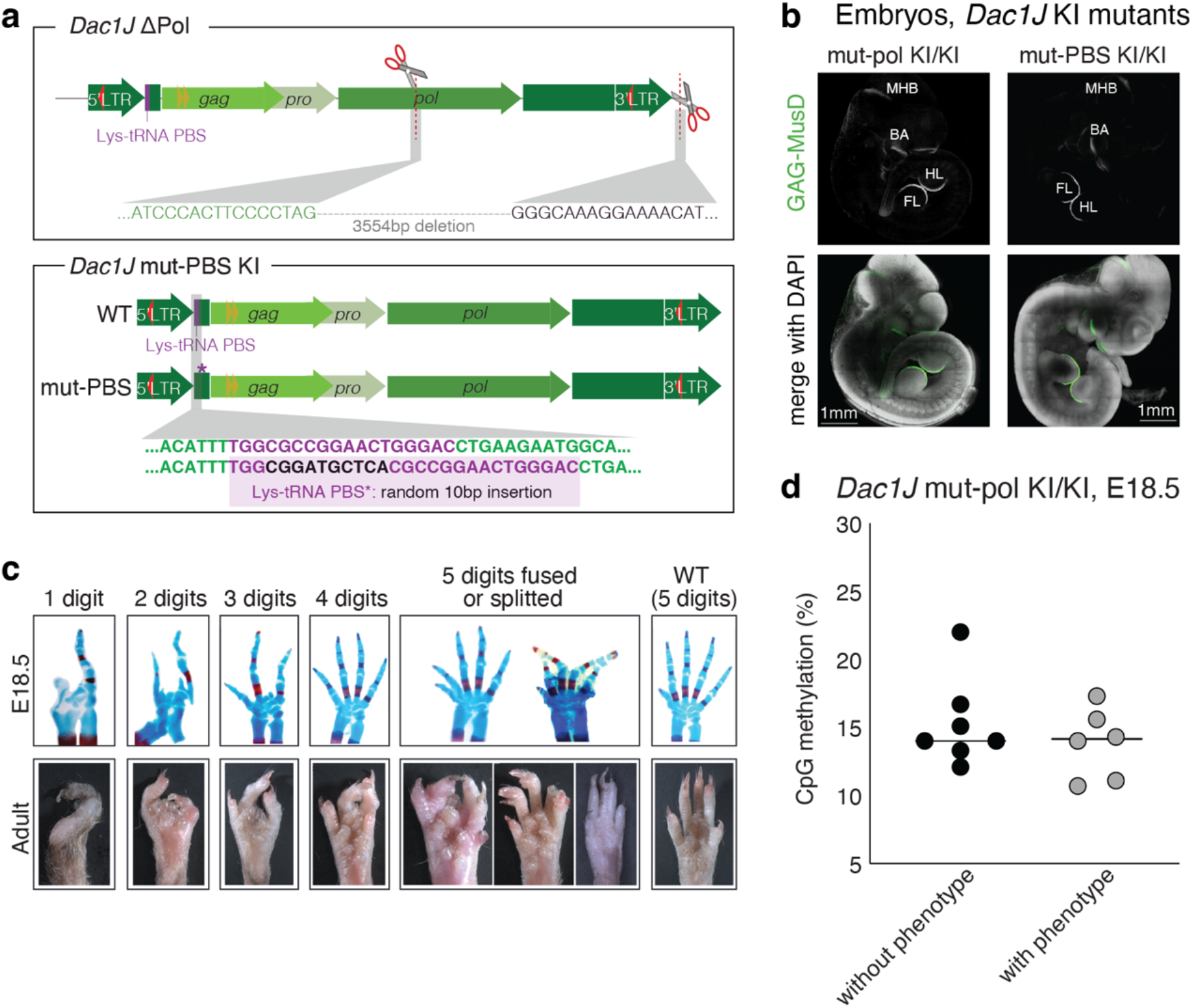
Knock-ins of MusD-*Dac1J* carrying pol and PBS mutation. **a**, Detail of the 3554bp deletion of the Pol gene and 3’ sequence (top) and of the 10bp insertion of random DNA in the *Dac1J*-mutPBS knock-in (bottom). In the Δ/ΔPol-*Dac1J* AER cells, Gag and Pro viral-like proteins exist in the AER cells and assemble into a VLP lacking reverse transcriptase. This incomplete VLP can neither replicate ETns, nor undergo tRNA primed reverse transcriptase of its RNA. In the *Dac1J*-mutPBS knock-in, the 10bp insertion in the Lys tRNA primer binding site (PBS) leads to a scrambled PBS. In the *Dac1J*-mutPBS AER cells, Gag, Pro, and Pol retroviral-like proteins are translated into the cytoplasm and assembled into VLP. The intact Pol allows possible replication of ETn elements but the MusD VLPs are unable to undergo tRNA-primed reverse transcription. **b**, anti-GAG-MusD whole-mount immuno-fluorescence on E10.5 Δ/ΔPol-*Dac1J* (left), mut-pol KI/KI (middle), and mut-PBS KI/KI (right) whole embryos. Scale bars 1mm. At least *n*=3 biological replicates were confirmed. FL, forelimb; HL, hindlimb; BA, branchial arches; MHB, mindbrain-hindbrain boundary. **c**, Representation of E18.5 skeletal analysis and adult limbs illustrating the 6 different observed phenotypes indicated in **Fig. 5d. d**, CpG DNA methylation at the 5’LTR promoter of the *Dac1J* mut-pol insertion measured by pyrosequencing in *n*=7 E18.5 embryos without a phenotype and *n*=6 E18.5 embryos with a phenotype. Each dot represents the average percentage of DNA methylation from 7 CpGs. Overall, no significant difference in average 5’LTR DNA methylation is observed between the animals with or without a phenotype.

**Extended data Fig. 9.**
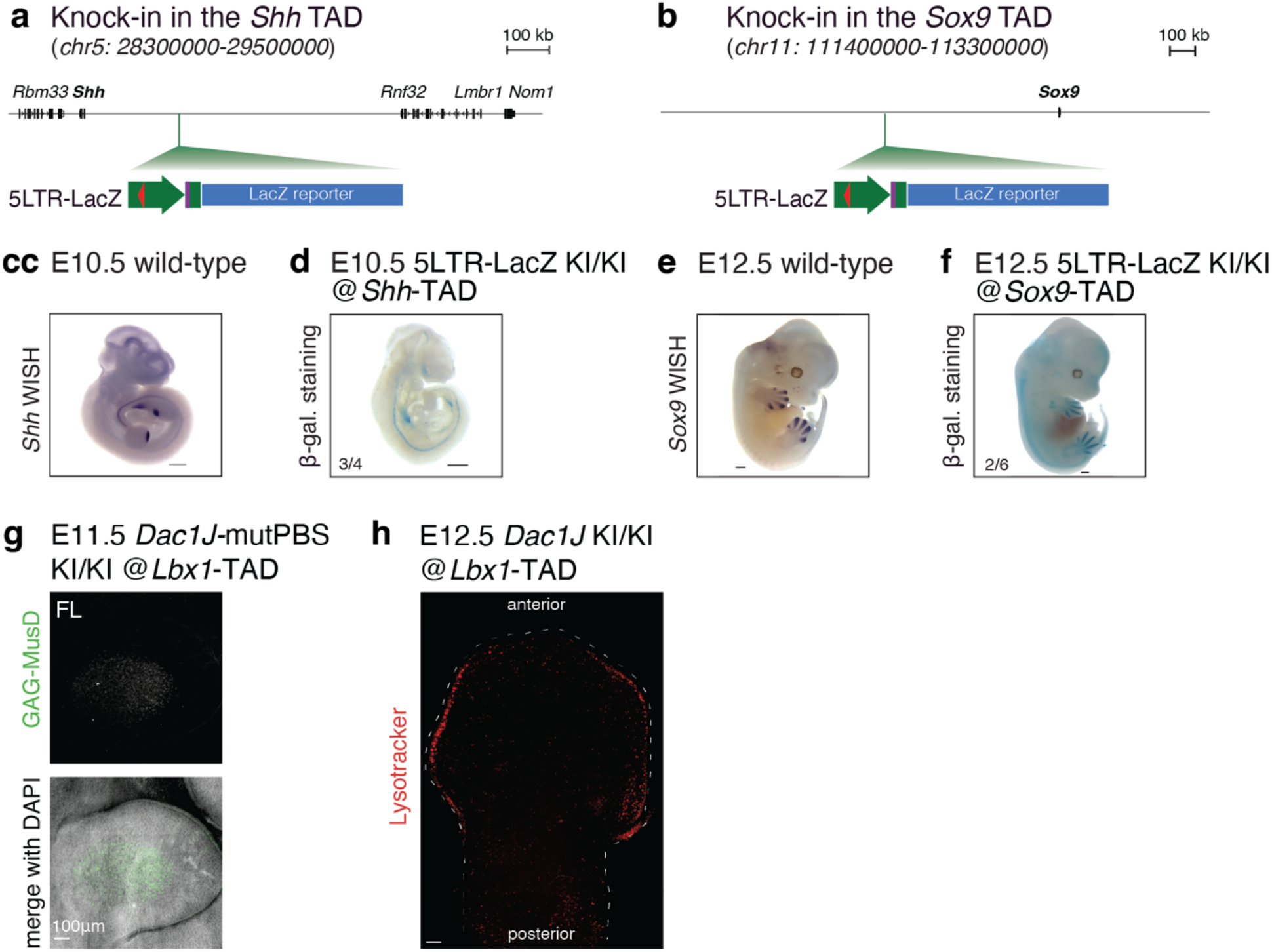
MusD-*Dac1J* adopts the expression of genes surrounding its insertion. **a-b** Schematic representation of the *Dac1J* -5LTR-LacZ knock-in at the *Shh* (**a**) and *Sox9* (**b**) locus. **c**, whole mount in situ hybridization of *Shh* from E10.5 wild-type embryo. Scale bar 500um. **d**, beta-galactosidase staining on E10.0 *Dac1J*-5LTR-LacZ (*Shh* knock-in) showing whole embryos. Scale bars 500um, n= 3/4 embryos show similar staining. **e**, whole mount in situ hybridization of *Sox9* from E12.5 wild-type embryo. Scale bar 500um. **f**, beta-galactosidase staining on E12.5 *Dac1J*-5LTR-LacZ (*Sox9* knock-in) showing whole embryos. Scale bars 500um, *n*= 3/4 embryos show similar staining. **g**, anti-GAG-MusD whole-mount immuno-fluorescence on E11.5 *Dac1J*-mutPBS KI/KI (*Lbx1* knock-in) showing forelimb staining. Scale bars 100um. At least *n*=3 embryos were tested. **h**, Lysotracker staining of an E12.5 *Dac1J* KI/KI (*Lbx1* knock-in) forelimb. Expected AER staining for that stage is observed but no muscle cell progenitor staining is detected. *n*=3 embryos were analyzed.

